# Range wide analysis of genetic diversity and structure gives insights into *Rosa gallica* L. evolutionary history

**DOI:** 10.64898/2026.08.05.742742

**Authors:** Clovis Pawula, Jérémy Clotault, Olivier Lepais, Annie Chastellier, Eduardo Javier Ordoñez Trejo, Tatiana Thouroude, Silvia Assini, Ladislav Bakay, László Bartha, Jože Bavcon, Jocelyne Cambecèdes, Jordane Cordier, Anna Cwener, Zygmunt Dajdok, Pavel Dřevojan, Jérôme Garcia, Jasmin Grahić, Adam Kapler, Viktor Kerényi-Nagy, Almira Konjić, Grzegorz Łazarski, Nicolas Leblond, Alexander Mrkvicka, Karel Nepraš, Halyna Oliiar, Marziano Pascale, Ivan Pejić, Renata Piwowarczyk, Blanka Ravnjak, Per Harald Salvesen, Veronica Sărăţeanu, Ivan Schanzer, Adriano Soldano, Elena Tofan-Dorofeev, Nikola Tomljenović, Karolina Wiśniewska, Mateusz Wolanin, Valéry Malécot, Agnès Grapin, Alix Pernet

**Affiliations:** Univ Angers, Institut Agro, INRAE, IRHS, SFR QUASAV, F-49000 Angers, France; Univ. Bordeaux, INRAE, BIOGECO, F-33610 Cestas, France; Univ. Pavia, Dept. of Earth and Environmental Sciences, I-27100 Pavia, Italy; Slovak Agricultural University, Institute of Landscape Architecture, Tulipánová 7, Nitra 94901, Slovakia; Molecular Biology Center, Interdisciplinary Research Institute on Bio-Nano-Sciences, Babes, -Bolyai University, 42 August Treboniu Laurean Street, 400271 Cluj-Napoca, Romania; University Botanic Gardens Ljubljana, Biotechnical faculty, Ižanska cesta 15, 1000 Ljubljana; Conservatoire botanique national des Pyrénées et de Midi-Pyrénées, BP70315, F-65200 Bagnères de Bigorre, France; Conservatoire botanique national du Bassin Parisien, Muséum national d’Histoire naturelle, Orléans, France; Botanical Garden, Maria Curie-Skłodowska University, Lublin, Poland; Department of Botany, Faculty of Biological Sciences, University of Wrocław, Kanonia 6/8, PL-50-328 Wrocław, Poland; Department of Botany and Zoology, Faculty of Science, Masaryk University, Kotlářská 2, 611 37 Brno, Czech Republic; Faculty of Agriculture and Food Sciences, University of Sarajevo, Zmaja od Bosne 8, 71000 Sarajevo, Bosnia and Herzegovina; PAS Botanical Garden – Center for Biological Diversity Conservation in Powsin, Prawdziwka 2, 02-973 Warsaw 76, Poland; 1041 - Budapest, Görgey Artúr utca 12; Institute of Biological Sciences, Cardinal Stefan Wyszyński University, Wóycickiego 1/3, 01-938 Warsaw, Poland; Conservatoire botanique national Sud-Atlantique, F-33980 Audenge, France; Perchtoldsdorf, Austria; Department of Preschool and Primary Education, Faculty of Education, Jan Evangelista Purkyně University in Ústí nad Labem, České mládeže 8, 400 01 Ústí nad Labem, Czech Republic; Medobory Nature Reserve, 21, Mitskevych str., Hrymailiv, Ternopil Region, Ukraine, 48210; Roccavione, Italy; University of Zagreb Faculty of Agriculture - Svetošimunska 25, 10000 Zagreb, Croatia; Center for Research and Conservation of Biodiversity, Department of Environmental Biology, Institute of Biology, Jan Kochanowski University, Uniwersytecka 7, PL-25-406 Kielce, Poland; Department of Biological Sciences, University of Bergen, Bergen, Norway; University of Life Sciences “King Mihai I” from Timis, oara, Agriculture Faculty, 300645 Timis, oara, Romania; Tsitsin Main Botanical Garden, Russian Academy of Sciences, Moscow, 127276 Russia; Vercelli, Italy; National Botanical Garden Institute, Moldova State University; Agronomy School Zagreb - Ul. Gjure Prejca 2, 10040 Zagreb, Croatia; Faculty of Biology and Nature Protection, Rzeszów University, Zelwerowicza 4, 35-601 Rzeszów, Poland

**Keywords:** Population genetics, Crop Wild Relatives, Polyploids, Interspecific Hybrid, SSRseq

## Abstract

*Rosa gallica* L., the French rose, is a perennial, tetraploid, heterozygous species that naturally propagates by seed and sucker. It occurs in the wild, primarily in Europe, and also exists as cultivated varieties. *R. gallica* cultivars were extensively bred and cultivated in France at the beginning of the 19th century. Although several hypotheses have been proposed regarding the species expansion based on historical records, none have been assessed using molecular data. Indeed, its genetic diversity has so far been investigated only at local or regional scales, hindering the identification of the evolutionary factors shaping its present-day distribution.

Using 29 sequenced microsatellites, we genotyped a comprehensive sample of 1618 individuals, including wild *R. gallica* from 219 sites across the species range, rose cultivars, and specimens from other *Rosa* species. We then detected clonal lineages and characterized the range-wide genetic diversity and structure, aiming to disentangle the roles of natural and human factors in shaping the distribution of *R. gallica*, with particular focus on France.

French diversity appears particularly structured compared to the rest of the range, suggesting multiple origins within France. Populations in South Alps, Central Eastern Europe, and Eastern France appear to have recolonized naturally from a single southern glacial refugium. In contrast, populations in the western part of France likely resulted from more recent natural or human-mediated dispersal. Finally, clonal lineages containing both wild and cultivated individuals were predominantly found in France, highlighting the role of human-mediated dispersal in 28 of the 98 French sites studied.

These findings show that the present-day natural range of *R. gallica* was shaped primarily by post-glacial recolonization, but also reveal a contribution of human activities to its recent dispersal, particularly in France, where cultivated varieties were intensively bred and exchanged.

## Introduction

The distribution of a plant species results from various biological, historical, and evolutionary factors such as dispersal dynamics, climate modifications, human activities, or hybridization (Ali, 2020; Bullock et al., 2018, 2017). Among these, human activities profoundly impact plant distribution patterns, primarily through species dispersal and environmental alterations (Bullock et al., 2018). Horticultural ornamental species demographic history may have been shaped by both natural and human dispersions as these have been extensively hybridized and propagated by humans. Moreover, these species often possess specific traits such as hardiness and rapid vegetative growth that enhance their survival in the wild (Maurel et al., 2016; van Kleunen et al., 2018). In addition, perennial ornamental plants can persist for decades or even centuries, making them valuable subjects for studying long-term effects of human actions on plant populations and biodiversity.

Roses have been cultivated and exchanged throughout the world for thousands of years. Human love for roses began during the Neolithic period and has endured through various ages and across cultures (Nybom, 2009; Tomljenović and Pejić, 2018; van Kleunen et al., 2018). Roses are appreciated for their beauty, fragrance, nutritional qualities and medicinal properties (Hegde et al., 2022; Nybom, 2009). Cultivated roses have been obtained from 8 to 15 species of the 150 to 200 recognized diploid or polyploid species in the genus *Rosa* (Smulders et al., 2011; Wissemann, 2003, 2017; Wylie, 1954). Roses are classified using two different classifications, the botanical one (Wissemann, 2017) and the horticultural one (Cairns, 2003). The botanical classification distributes wild rose species into four subgenera, the largest subgenus (the subgenus *Rosa*) being divided into 11 sections. Debray et al. (2022) have deeply examined this mostly morphology-based classification in the light of genomic data and suggest to merge certain sections. The horticultural classification of the roses was proposed by the American Rose Society and then adopted by the World Federation of Rose Societies. It divides the roses into three classes, wild and botanical roses, Old Garden Roses (OGR) bred before 1867, and modern roses bred after this year. OGR and modern roses are subdivided into horticultural groups that include cultivars with similar traits (Cairns, 2003).

*Rosa gallica* L., also known as the ‘French Rose’, is one of the main ancestors of the European OGR and of the oil-bearing roses. The OGR cultivars the most closely related to *R. gallica* would belong to the horticultural group Hybrid Gallica while other related cultivars would be found into the Damask, Centifolia, Alba, Portland, Moss and Bourbon horticultural groups (Roberts et al., 2003). *R. gallica* has been cultivated since antiquity, but most cultivars were bred by French rose breeders in the early 19th century, who produced over two thousand cultivars (Joyaux, 1998). Currently, wild *R. gallica* are found mainly in Europe, with the natural range extending from the Middle East to France (Ercisli, 2005; Khapugin et al., 2021; Klášterský, 1968). These wild populations are protected in France (https://www.legifrance.gouv.fr), Poland (Wójcik and Ziaja, 2022) and in Czechia (Grulich, 2012). According to Fougère-Danezan et al. (2015), *R. gallica* have originated less than 10 millions years ago, certainly during the Pliocene or at the beginning of the Pleistocene. *Rosa gallica* is a small shrub with the ability to produce root suckers (Boulenger, 1932) generating clonal individuals. The species is a segmental allotetraploid that may have an heredity in the disomic to polysomic continuum (Fernández-Romero et al., 2001). Finally, as commonly described in the genus, *R. gallica* can naturally hybridize with other *Rosa* spp. (Fedorova et al., 2010).

The origins of the French and European *R. gallica* populations are unclear. As the *R. gallica* species appeared before the last glacial maximum (18-25 kyr BP), one hypothesis would involve a natural recolonization of Europe after this period from one or several Southern refugia, as described for other perennial species (Ali, 2020; Tzedakis et al., 2013). Other hypotheses, implicating human-mediated dispersal, have been proposed on the basis of historical reports. One suggested that *R. gallica* might have colonized Europe after escaping from Roman fields (Debray, 2020). Other authors hypothesized that the species may have been introduced to Europe from the Middle East since the 13th century (Saakov, 1965). For the most Western part of the range (West of France and Spain), Klášterský (1968) hypothesized that *R. gallica* populations in these areas may have originated from secondary introductions. In the same manner, Boulenger (1932) indicates that *R. gallica* has been widely cultivated in France and may have escaped from the gardens in some localities, this may result in the coexistence of indigenous populations and individuals of cultivated origin.

Explorations of *R. gallica* diversity have been performed only at limited scales, using either morphological or genetic markers. In Poland, palynologists have identified two different groups of pollen shapes among Polish *R. gallica* (Wrońska-Pilarek and Boratyńska, 2005). The most extensive study, in terms of geographic area, was limited to around forty samples from Ukraine, Hungary, Romania, and Greece. The authors used Inter Simple Sequence Repeats and revealed a clear spatial genetic structure (Fedorova, 2014; Fedorova et al., 2010). De Cock (2008) compared populations from South East France to a German one using AFLP, and found the German individuals to be differentiated from the French ones. At the France scale, preliminary works revealed a structure between wild and cultivated *R. gallica*. Additionally, wild French specimens that were collected in geographically close sites were genetically similar (Pernet et al., 2009; Thouroude et al., 2012). Another study focused on South East France and used morphological and RAPD data to reveal a high genetic diversity (Reynders-Aloisi et al., 2000).

In this study, our main objective is to better understand the origins of *R. gallica* populations across its range, particularly in France. To achieve this, we undertook extensive sampling of rose specimens, including *R. gallica* collected from a large part of its natural range, other *Rosa* species, and cultivated roses. We genotyped these samples using a recently designed SSR sequence-based genotyping system (Pawula et al., 2023). Subsequently, we performed clonal lineages and interspecific hybrid detection based on Euclidean distances and on custom dissimilarities, respectively. We then obtained a refined dataset, including only non-hybrid *R. gallica* specimens, to answer the following questions: (1) Do French populations and those from other countries share a common demographic history? (2) Are the hypotheses on *R. gallica* demographic history, found in literature, supported by present-day genetic diversity and structure? (3) Is there evidence of human-mediated dispersal, as suggested in historical references?

## Materials and methods

### Sampling

Wild sites of *R. gallica* are often composed of a few continuous plots, each covering few to dozens square meters. Due to the production of root suckers, each plot generally contains a few clonal individuals. Therefore, a site may regroup only a low number of genets (i.e. different clonal lineages, a genet being a group of individuals related to each other by asexual reproduction). In the present study, we use “site” instead of “population” to refer, for wild specimens, to the collecting place. Indeed, according to Waples and Gaggiotti (2006), “a population is group of individuals of the same species living in close enough proximity that any member of the group can potentially mate with any other member”. The definition implies the notion of reproductive cohesion that is difficult to determine *a priori*, based on the sampling location and on the geographical distances between individuals (Waples and Gaggiotti, 2006). Therefore we preferred to use “site” that does not imply such reproductive cohesion but only geographical co-occurrence. Arbitrarily, individuals were considered belonging to two different sites when more than one kilometer separated all individuals from each site. Given the low expected number of genets per site, the number of collecting sites was maximized lowering the number of individuals collected per site. In total, leaf samples were collected on 905 *R. gallica* individuals from 223 sites in 15 countries (Fig. 1). Among them, 485 individuals were collected from 102 French sites. The number of individuals collected and genotyped from each site ranged from one to thirteen, with a median of four. When possible, we tried to collect at least four individuals per site by collecting at least one individual from each continuous plot of plants present on the site. Therefore more individuals were collected in sites harboring many plots. In addition, samples of 402 rosebushes suspected to be related to *R. gallica* were gathered. These were either botanical individuals attributed to *R. gallica* (e.g. *R. gallica* var. *incarnata*) or cultivated individuals belonging to the Hybrid Gallica and other OGR groups related to *R. gallica*: the Damask, Centifolia, Portland, Alba, Bourbon or Moss groups. Botanical individuals of *R. gallica* are plants, of suspected wild origins, preserved in rose gardens named with latin epithet following the botanical code (Turland et al., 2018) while cultivated individual belongs to horticultural groups of OGR named with fancy names following the cultivated plant code (Brickell et al., 2016, e.g. ‘Rosiers des parfumeurs’). To be able to detect species mis-identification or *R. gallica* interspecific hybrids (either wild or cultivated) with other *Rosa* spp., we gathered 259 more individuals belonging to other species of the genus. These were wild, botanical, or cultivated specimens not related to *R. gallica*, chosen to represent the diversity of the genus *Rosa* according to findings of Debray et al. (2022). Finally, we included 10 full-sibling and 10 half-sibling individuals to evaluate the resolution of clonal lineages detection. Samples metadata and details are available in Table S1.

**Figure 1.**
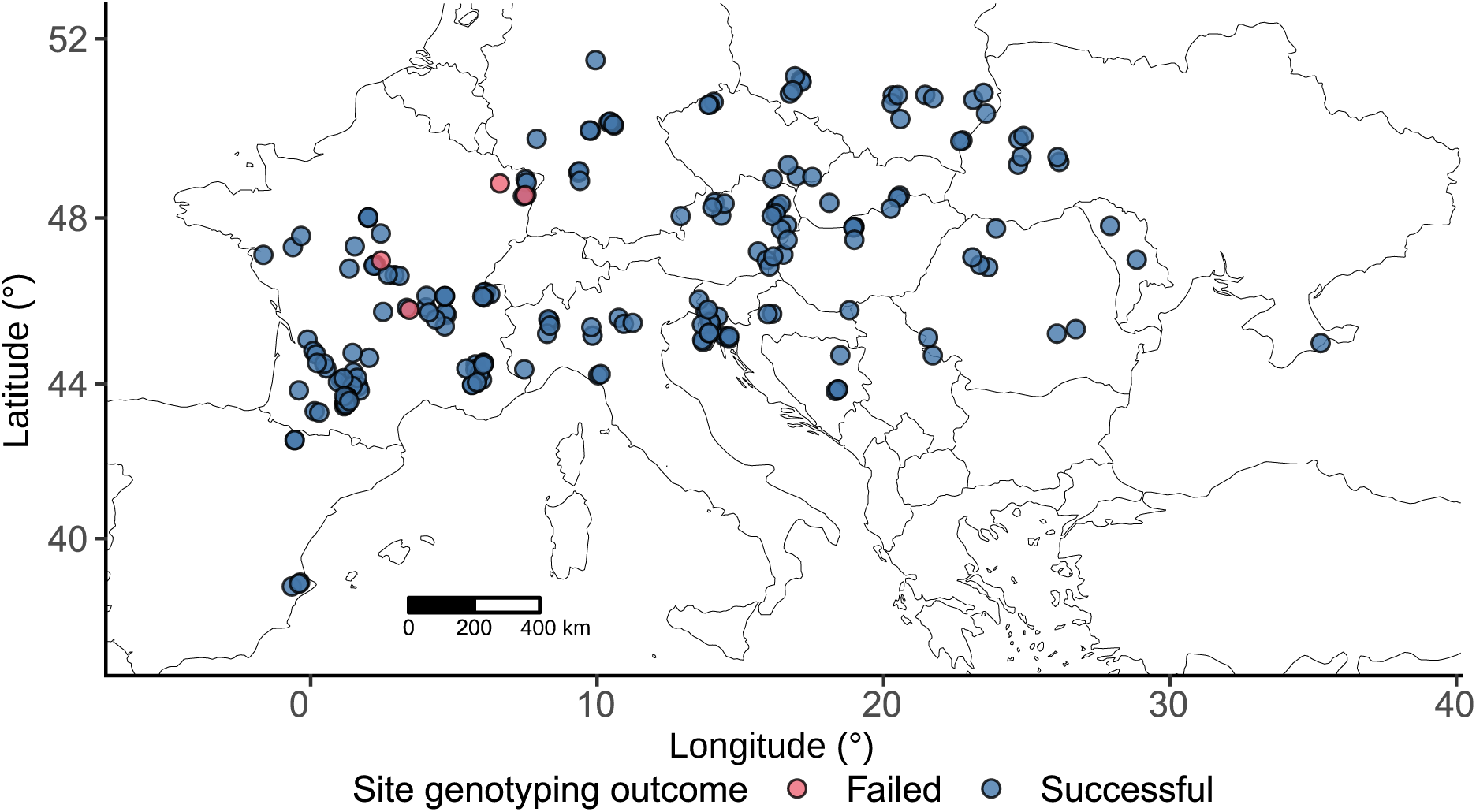
Wild sampling sites location and genotyping outcomes. Sites are colored according to whether at least one individual was successfully genotyped (blue, 219 sites) or none (pink, four sites).

### DNA processing and genotyping

As the samples were either gathered from former projects, from the RosePom Biological Resources Center (Angers, France) or especially for this project by different collectors, there was variability on their nature. They were either freeze dried, silica dried or already in the form of extracted DNA. A few samples were obtained from herbarium specimens. Formerly extracted DNA were isolated using the Qiagen 96 Plant Kit (Qiagen, Hilden, Germany), the NucleoSpin® 96 Plant II Core Kit (Macherey-Nagel, Düren, Germany) or the protocol described in Debray et al. (2022). DNA extracted for this project were obtained using a custom protocol inspired from Keb-Llanes et al. (2002), Inglis et al. (2018) and Anderson et al. (2018). More precisely, 20-30 mg of freeze dried or silica dried leaves samples were put in wells of a 2.2 ml 96 wells plate, for herbarium specimens already dried leaves were used. Samples were ground in a ball mill with stainless steel balls for 3-6 minutes at 1500 Hz. Then, they were pre-washed according to the sorbitol pre-wash step described in Inglis et al. (2018). Lysis step was performed using the CTAB-SDS based methods described in Keb-Llanes et al. (2002) with modifications. These include addition of 0.5 µl per sample of RNAse A 20 mg.ml*^−^*^1^ to the EBA buffer and addition of 2 µl per sample of Proteinase K 20 mg.ml*^−^*^1^ to the EBB buffer. Extraction buffer volumes were also reduced to 200 µl EBA, 600 µl EBB and 70 µl of SDS 20% per well. Incubation was performed for 1 hour at 65 °C while shaking every 15 minutes. Precipitation step was also done according to Keb-Llanes et al. (2002). For this step, the centrifugation speed was set to 5800 g and the obtained clean DNA pellets were resuspended in 50 µl of TE buffer pH 8 (10 mM Tris pH 8, 0.1 mM EDTA pH 8). DNA quantities were then assessed using the DNA-specific fluorescent Hoechst 33258 and a microplate reader (FLUOstar Omega, BMG Labtech, Ortenberg, Germany). Subsequently, the DNAs were normalized to 10 ng.µl*^−^*^1^.

In total 1710 samples were genotyped, including 1586 unique individuals plus technical replicates (Table S2). The sequencing was done in five different runs including a pilot run where 95 samples (including diploids and tetraploids), chosen to represent the genus diversity, were genotyped twice. Individuals were genotyped using 60 primer pairs targeting nuclear SSRs (Table S3). Primer pair design is described in our previous work (Pawula et al., 2023). The Polymerase Chain Reaction (PCR) process, library preparation and sequencing experiment were carried out at the Genome Transcriptome Facility of Bordeaux, following the method outlined in Lepais et al. (2020), with a few modifications. Instead of performing the multiplexed locus-specific PCR in two separate steps, it was done in a single step. Additionally, the number of PCR cycles was reduced to 24 cycles from the original 35 cycles. Once the individuals were pooled, the sequencing step was performed on an Illumina iSeq100 using a 2 *×* 150 bp paired-end sequencing kit.

### Allele calling

The raw read sequences were sorted using the Illumina barcodes, and the adaptors were eliminated using the software provided by the sequencer platform. Paired-end reads were merged using BBmerge, with a minimum overlap length of 20 bp (Bushnell et al., 2017). Additionally, merged reads shorter than 65 bp were discarded for further analyses. Raw read sequences were converted into the final microhaplotypes using the pipeline based on the FDSTools v1.2 analysis toolkit (Hoogenboom et al., 2017) designed by Lepais et al. (2020). The detected alleles were coded using an arbitrary three-digit code.

Several parameters were needed to run the pipeline (Hoogenboom et al., 2017). Values for these parameters have a direct impact on the calling success (allelic errors and missing data, Lepais et al., 2020). Therefore, several sets of parameters were tested for each locus. To evaluate the parameter sets, we used the data of known diploid and tetraploid individuals from the pilot sequencing run that were successfully sequenced twice (87 individuals, Table S2). Since one of the parameters corresponded to the ploidy level, the allelic error rate and the missing data rate were separately calculated for the 42 diploid and 45 tetraploid individuals. For a given locus, the allelic error rate was calculated using the formula:

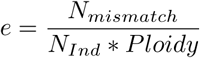

with *N_mismatch_*being the number of allelic mismatches over all comparisons and *N_Ind_* being the number of individuals compared with their replicates. The expression “*N_Ind_ ∗ Ploidy*” is therefore equal to the number of alleles compared. One of the parameters was the analytical strategy. The two possible strategies were either the *FullLength* or the *RepeatFocused* methods. In the *FullLength* strategy, the calling considered all the variations located between the two primers whereas, in the *RepeatFocused* strategy, flanking sequences were ignored and only the sequence variations in the repeat motif were considered. Focusing on the repeat motif allowed to reduce the allelic error rates, by not taking into account error prone flanking sequences. For each locus, the parameter set producing the lowest allelic error and missing data rates, for both diploids and tetraploids, was selected. Final parameters are available in Table S4. These parameters were then assumed to be optimized whatever the ploidy level. To remove loci that were obviously paralogous, alleles were called by allowing up to 20 alleles at a locus per individual. Then, we implemented the outlier detection method of Rosner available in the EnvStats R package (Millard, 2013; Rosner, 1975) to identify loci with significantly more alleles than others and thus potentially paralogous. These were removed for further analyses.

The ploidy parameter is crucial in the calling process. Setting it lower than the actual ploidy level increases the number of loci with missing data because the pipeline classifies loci with a higher number of alleles than the ploidy parameter as missing. Inversely, setting the ploidy level higher than the true ploidy level could relax the error detection and then increase the number of errors (Hoogenboom et al., 2017). Therefore, after the setting of parameters of allele calling by locus, we determined the ploidy level of the individuals. Due to the heterogeneous conditioning of the samples, we were unable to perform chromosome counting or flow cytometry. Several methods have been proposed to aid ploidy determination in polyploids using codominant data. Simplest tools used the maximum number of identical-by-state alleles found for an individual to set its ploidy level (Clark and Jasieniuk, 2011; Meirmans, 2020). However, this method requires a careful examination of each sample and is sensible to allelic errors that can easily increase the number of different alleles at a locus. Huang et al. (2019) developed a method based on an iterative process coupled with allele frequency estimation to determine the ploidy level of individuals having a polysomic inheritance in a population assuming panmixia within each ploidy level. Unfortunately, this method was not suitable for our data due to the likely absence of a polysomic inheritance mode (Bourke et al., 2017; Fernández-Romero et al., 2001; Lunerová et al., 2020). Additionally, we expected departure from panmixia in our sample given the high number of different species represented. Therefore, to solve this statistical classification problem, we implemented a supervised machine learning method known as Random Forest using the R package caret v 6.0.93 (Kuhn and Max, 2008). As a supervised algorithm, the Random Forest model requires a training set. This means that we needed a group of individuals with known ploidy levels that represent all potential ploidy levels in our sample. We established a group of 194 individuals with high confidence in their ploidy level based on measurements done in other studies or previously in our lab (Debray et al., 2022; Fernandez-Romero et al., 2009; Liorzou et al., 2016; Roberts et al., 2009). The number of triploids and pentaploids included in the present study was expected to be low. In accordance, few of them were present in the training dataset, which limited high-quality predictions for these two ploidy levels. To mitigate this, we decided to limit the prediction to the three main ploidy classes anticipated in our sample: diploids, tetraploids, and hexaploids. To achieve this, we categorized the reference samples into these three classes by merging triploids with tetraploids and pentaploids with hexaploids. To conduct the training and prediction, individuals were called allowing up to 20 alleles at each locus. We calculated the number of alleles at each locus and trained the model using the reference groups. The model accuracy was estimated using the out-of-bags estimates. We then utilized this model to predict the ploidy level of the other individuals in our sample. The final allele calling was done using the predicted ploidy level value as the ploidy parameter of the FDSTools-based pipeline.

Retrieving the allele dosage in diploids is straightforward as it can be deduced from the number of identical by state alleles. For polyploids, the number of alleles can only provide information about the allele dosage when either it is equal to one (only one allele is present at a locus) or the number of different alleles equals the ploidy level. The non straightforward retrieval of allele dosage is known as genotype ambiguity. To address this ambiguity and estimate the allele dosage, we utilized the method published by Cui et al. (2022). This method uses read depth and a correction for preferential amplification to infer the allele dosage. We performed the computation using our own R implementation of this method.

To minimize the impact of missing data on our sample, we first removed loci and samples with more than 40% of missing data. After that, a custom R script (Eveilleau and Pernet pers.com. modified by Pawula) was used to optimize the handling of missing data by selectively removing either loci or individuals. The aim was to retain the maximum amount of data in the dataset while ensuring that each pair of individuals can be compared using at least half of the loci.

### Marker evaluation

At this step, the dataset included 1618 samples successfully genotyped (including known repeated individuals) and 29 loci. Marker information was recorded on this dataset, designated hereafter as the *Rosa* spp. dataset (Fig. 2). In detail, the missing data rates, the observed number of alleles considering Whole Amplicon Information (WAI, i.e. sequence identity), the observed number of alleles considering only amplicon length, the number of rare alleles (allele frequency lower than 0.01), and the Polymorphism Information Content (PIC) were determined. The PIC was calculated using the function PIC available in the R package Polysat v1.7.7 (Clark and Jasieniuk, 2011).

**Figure 2.**
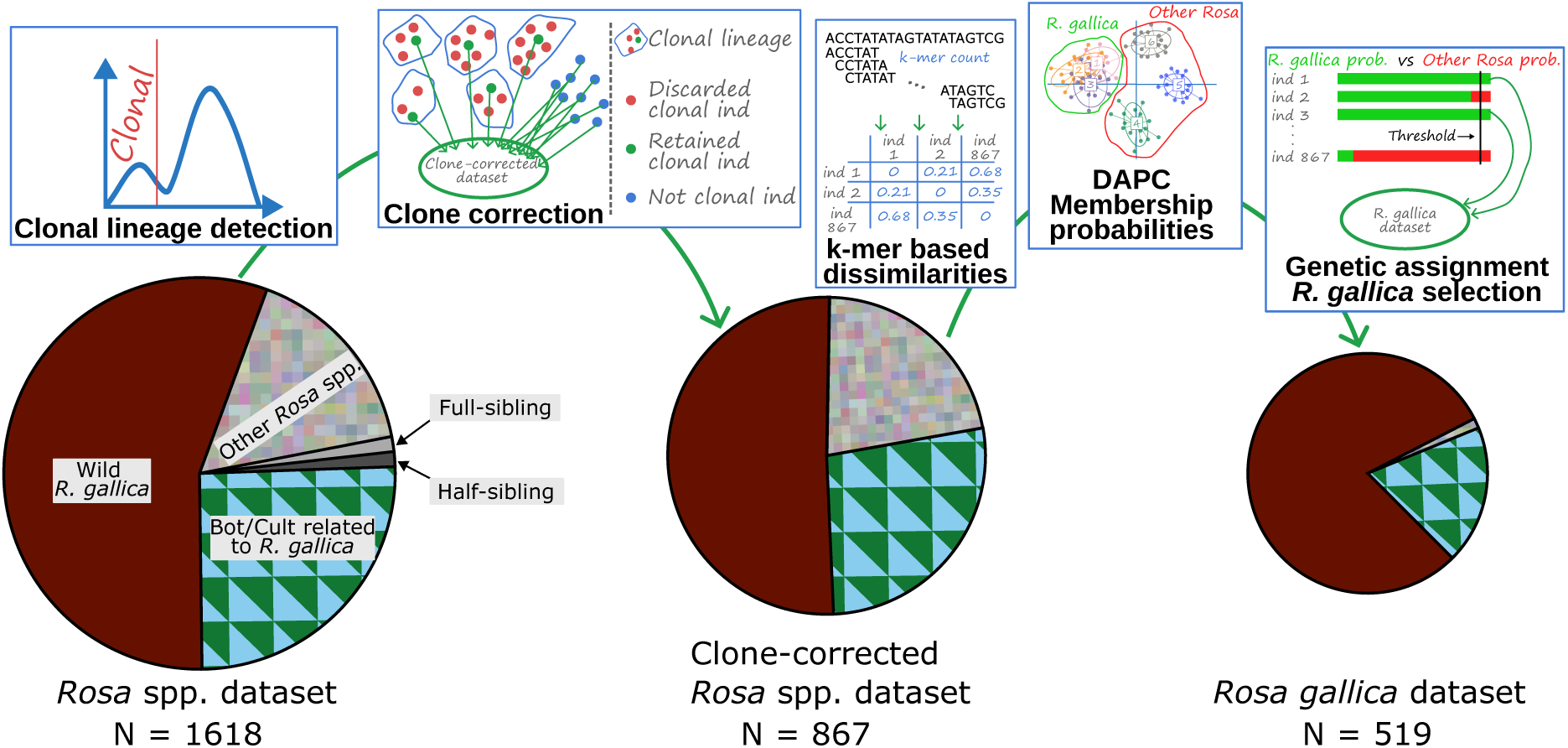
Workflow of individual filtering. Of the 1710 samples, 1618 were successfully genotyped, constituting the *Rosa spp.* dataset. This dataset included individuals initially identified as wild *R. gallica*, those from other *Rosa* species, and botanical or cultivated individuals presumably related to *R. gallica*. Additionally, known full-sibling and half-sibling samples were introduced to evaluate the false positive rate of clonal lineage detection. Clonal lineage detection was then performed, and the dataset was clone-corrected by retaining only one individual per clonal lineage and all non-clonal individuals. Known full-siblings and half-siblings were removed at this stage. This refined dataset was designated as the clone-corrected *Rosa spp.* dataset. Genetic assignment was subsequently conducted to exclusively select *R. gallica* individuals. This final dataset, called the *R. gallica* dataset, contained individuals initially collected as wild *R. gallica* and botanical or cultivated individuals. Remarkably, seven individuals, initially considered as belonging to other *Rosa* species, were found to be *R. gallica* specimens (Table S2), they were included in the *R. gallica* dataset.

An important metric for assessing marker quality is the extent of null alleles. Null alleles occur when the genotyping method fails to identify the presence of true alleles. For a diploid individual, this makes the loci with a null allele to appear as missing data if it is homozygous for the null allele or to appear as homozygous in case of it has an heterozygous genotype with a detectable allele and a null allele. Consequently, null alleles induce an increase in F*_is_* statistics, suggesting a departure from panmixia due to an excess of homozygotes (De Meeûs, 2018). However, null alleles are not the only cause of an apparent excess of homozygotes as some other causes are Wahlund effect and selfing (Waples, 2015). Various methods are available to estimate allele frequencies in the presence of null alleles for diploids under the assumption of Hardy-Weinberg equilibrium (Dąbrowski et al., 2015). For polyploids, dosage uncertainty, inheritance mode and double reduction make null allele estimation even more complicated. Huang et al. (2020) adapted the methods of Kalinowski and Taper (2006) to estimate allele frequencies in the presence of null alleles for autopolyploids. However, their method was unsuitable for our data because it requires groups of individuals in Hardy-Weinberg equilibrium and uses models of polysomic inheritance. De Meeûs (2018) proposed a method to determine if null alleles are impacting the data in non-equilibrium populations. Despite being tested only on diploids, as this method is based on F-statistics and missing data rates, it should be applicable to tetraploids. Therefore, this method was used to ascertain if null alleles were affecting our dataset. In detail, the clusters inferred by STRUCTURE at *K* = 8 (see below) with more than 20 non-admixed individuals were selected to form groups with reduced Wahlund effects. Polygene v1.6 was used to calculate F*_is_* and F*_st_* (Huang et al., 2020). Following the procedure proposed by De Meeûs (2018), the variation of F*_is_* and F*_st_* across loci, the Pearson’s correlation between F*_is_* and F*_st_*, the Pearson’s correlation between F*_is_* and missing data rates, and the differences between standard errors of F*_is_* and F*_st_* were examined. Pearson’s correlation coefficients were calculated using the R package ggpubr (Kassambara, 2023), while standard errors were determined using jackknifing over loci as done in De Meeûs (2018).

### Clonal lineages detection

Clonal lineages were detected using a threshold-based method, which has been adapted for polyploids and is available in the R package Polysat v1.7.7 (Clark and Jasieniuk, 2011). In details, pairwise Euclidean distances between individuals were computed based on intra-individual allele frequencies and using the R package factoextra v 1.0.7 (Kassambara and Mundt, 2020). The twice-genotyped individuals were used to establish a threshold to categorize pair of individuals as belonging to the same clonal lineage. In order to avoid any false-positive clonal relationship (i.e. two individual belonging to the same clonal lineage), the threshold was set to a conservative value. This corresponded to the maximal pairwise distance between the repeated individuals after the removal of outliers (i.e. distance values higher than the 1.5*×* interquartile range + third quartile of all pairwise distance between repeated individuals). Doing that, a threshold of 1.603 was used (Fig. S1). This threshold method has the advantage of permitting mutations and genotyping errors. However, it can also lead to a confusion between clonal and sibling individuals. To assess whether our resolution was strict enough to avoid this kind of confusion, we used 10 known full-sibling and 10 half-sibling individuals to verify whether they were incorrectly included in the same clonal lineage. These sib individuals were then removed from the dataset for further analyses. The clonal relationships between individuals from two sites or between wild and botanical or cultivated individuals were then visualized using the ggplot2 R package (Wickham, 2016). One individual per clonal lineage was retained for further analyses by selecting the most consensual with the less missing data as possible. This dataset is designated hereafter as the clone-corrected *Rosa* spp. dataset (Fig 2).

### Sequence based inter-individual dissimilarity

To take into account all the polymorphism types that could be retrieved from the marker sequences (repeat number variation, InDel and SNP), we developed a new measure of pairwise dissimilarity. The measure is a word-based alignment-free approach that uses the count of subsequences (i.e. k-mer) to estimate, for each locus, the evolutionary distance between the alleles. The procedure can be divided into three steps and is inspired from Kosman and Jokela (2019). First, we estimated the evolutionary distance for each pair of alleles at each locus. Second, we calculated dissimilarities between each pair of genotypes at each locus. Finally, we computed the dissimilarities between pairs of multilocus genotypes using all shared loci between each pair of individuals. These three steps are detailed below.

The first step was achieved using Alfpy v1.0.6 (Zielezinski et al., 2017) to perform k-mer counting and to calculate the dissimilarity between alleles at each locus. The k-mer length was set to six. We chose to use the Bray-Curtis dissimilarity measure as it takes into account the abundances of each k-mer and was shown to perform well in a benchmarking study (Zielezinski et al., 2019). For the second step, the dissimilarity between two genotypes *A* and *B* at a given locus *j* was determined in a very similar way as the one proposed by Kosman and Jokela (2019). We kept the nomenclature of these authors for consistency. First case: let consider individuals *A* and *B* are *q*-ploid, meaning that they respectively harbored the alleles < A_1_A_2_*…*A*_q_* > and < B_1_B_2_*…*B*_q_* > at the locus *j*. At the beginning of this step, all possible configurations for matching alleles from *A* to *B* were retrieved. For a given configuration, each allele appeared once and all alleles were present. The configuration was then composed of *q* allele pairs. The number of possible configurations was equal to *q*!. For diploid organisms, the number of possible configurations was 2. If *A* and *B* were diploid (*q* = 2) and respectively harbored the alleles *A*_1_, *A*_2_ and *B*_1_, *B*_2_ at the locus *j*, the possible configurations were *A*_1_*/B*_1_; *A*_2_*/B*_2_ or *A*_1_*/B*_2_; *A*_2_*/B*_1_. Then, the dissimilarity was calculated for each configuration using the allele Bray-Curtis pairwise dissimilarities computed at the previous step. The minimal value among these configurations was then taken as the dissimilarity between the two genotypes at the locus *j*. For *A* and *B*, this can be expressed as follows:

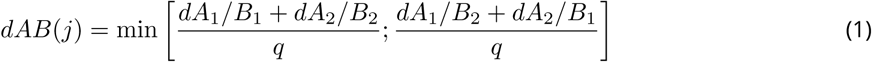

*dAB*(*j*) is the dissimilarity between genotypes *A* and *B* at the locus *j*. *dA_i_/B_k_* is the dissimilarity between allele *A_i_* and *B_k_* at the locus *j*.

Second case: let consider individuals with different ploidy levels. This requires additional steps. Our approach varied depending on the ploidy levels being compared. For a comparison between a diploid and a tetraploid, the diploid genotype was simply duplicated. For example, the heterozygote genotype *A* was considered as *A*_1_; *A*_2_; *A*_1_; *A*_2_. In the case of a diploid-hexaploid comparison, the diploid genotype was tripled. When comparing a tetraploid and a hexaploid, all possible hexaploid genotypes derived from the tetraploid genotype were determined. For instance, if a tetraploid genotype *D* had the genotype *D*_1_; *D*_2_; *D*_3_; *D*_3_, then the six possible hexaploid genotypes were:

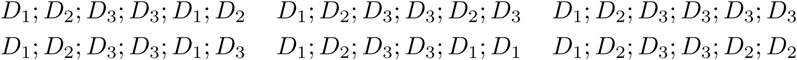

All the possible configurations were then determined based on these possible genotypes of *D*. As indicated in the equation (1), the smallest dissimilarity was retained as the dissimilarity between *D* and the hexaploid genotype at the locus *j*.

In the third step, to compute the final individual pairwise dissimilarities, the dissimilarities per locus were normalized by dividing all values by the maximum value in the dataset for each locus. Finally, dissimilarity between two individuals were computed by averaging their dissimilarities per locus.

### Mis-identification detection

We used the clone-corrected *Rosa* spp. dataset to select only *R. gallica* individuals for further analyses by detecting mis-identifications or putative hybrids. To do that, we first visualized the genetic structure within this dataset by performing a Principal Coordinates Analysis (PCoA) based on the pairwise k-mer dissimilarity matrix using the dudi.pco function from the R package ade4 v1.7.22 (Dray and Dufour, 2007). To compare this k-mer dissimilarity based method to a more conventional one, we also performed a Principal Component Analysis (PCA) based on intra-individual allele frequencies using the dudi.pca function from the R package ade4 (Dray and Dufour, 2007). Intra-allele frequencies were calculated based on the allele dosage, enabling the analysis of individual of various ploidy level using a PCA. For this PCA, missing data were turned as zero values.

After this step, a K-means clustering (R package adegenet v2.1.10, Jombart et al., 2010) was used to define groups of genetically similar individuals based on all the principal components of the PCoA. The clustering and determination of the most probable number of groups (*K*) were achieved by independently running the function find.clusters 100 times for each *K* from *K* = 1 to *K* = 30. We selected *K* = 12 as the most probable number of groups based on the Bayesian Information Criterion (Fig. S2). We then performed, on these principal components, a Discriminant Analysis of Principal Components (DAPC) to assign each individual a probability of belonging to each group. The function dapc.dudi was run on the first 11 principal components chosen using the *K*-1 criterion, as suggested by Thia (2023). We retained the first eight discriminant functions. Based on the scatter plot of the first two discriminant functions and on DAPC genetic group content, we selected the four groups out of the 12 that contained almost only individuals phenotypically attributed to *R. gallica* during the sampling process. This included a group, very close to the other three groups, containing a majority of cultivars belonging to the Hybrid Gallica horticultural group. By summing the membership probabilities for these four groups, we determined a probability for each individual to belong to the *R. gallica* species. Checking the distribution of the obtained probabilities (Fig. S3), we classified all the individuals having a probability higher than 0.99 as belonging to *R. gallica* for further analysis. This dataset is designated hereafter as the *R. gallica* dataset (Fig 2).

### Genetic structure and diversity of *R. gallica*

We investigated the population structure on the *R. gallica* dataset using different methods. First, we ran two multivariate analyses, one based on k-mer dissimilarities (PCoA) and one based on intra-individual allele frequencies (PCA) using the R package ade4 (Dray and Dufour, 2007). We performed these two multivariate analyses to not only investigate the *R. gallica* population structure but also to compare these two methods. Second, we used the hypothesis-driven bayesian clustering procedure implemented in the program STRUCTURE v2.3.4 (Pritchard et al., 2000). This model aims to define genetic groups while maximizing Hardy-Weinberg equilibrium inside each group. STRUCTURE parameters were selected following the recommendations of Wang (2017). Specifically, we used the ancestry model with admixture, while inferring the ALPHA parameters for each population (i.e. each STRUCTURE cluster), starting with an initial ALPHA value of 0.1. The inference was conducted using the correlated allele frequency model with a constant LAMBDA parameter. Based on the recommendations in the STRUCTURE manual and on *F_st_* plots, the burn-in period was set to 50 000 and the number of repetitions after burn-in to 300 000. The probability of the data under the model was estimated to identify optimal values of *K* groups. We ran the model with *K* values ranging from 1 to 20, with 10 independent runs per *K* value. The method of Evanno et al. (2005) was used to determine most probable *K* values. However, while this method is able to reveal large population structure, it generally performs poorly at detecting fine structure (Janes et al., 2017; Kalinowski, 2011; Puechmaille, 2016; Schwartz and McKelvey, 2009; Wang, 2019). Recently, Wang (2019) developed a parsimony estimator to detect optimal *K* values, implemented in KFinder v2.1. Wang (2019) demonstrated the robust performance of this estimator across various scenarios, including different population structure models, levels of differentiation, numbers of loci, and unbalanced sampling. Therefore, we used KFinder v2.1 to determine optimal *K* values aiming at revealing fine structure level. STRUCTURE inference can be stuck at local maxima, leading, for the same value of K, to independent runs sometimes returning different clustering. This situation may result in various modes of groupings, supported by a various number of independent runs. We used Pong v1.5 (Behr et al., 2016) to align the different runs, identify the modes, and visualize the ancestry proportion matrices (known as Q-matrices) in bar plots. Individual bars were clustered according to regional grouping. At the *K* determined by KFinder, the major mode was selected to calculate the mean ancestry proportions per site and to map these values at their GPS locations using the R packages pophelper v2.3.1 (Francis, 2017) and ggplot2 v3.4.1 (Wickham, 2016). Given the distribution of ancestry proportions (Fig. S4), we set 0.80 as the threshold to differentiate between admixed and non admixed individuals (Table S5). Finally, we used the STRUCTURE inferred allele frequency divergences among the genetic clusters at the major mode to draw an unrooted splits network using the lpnet R package v0.1.0 (Guo, 2023). The obtained nexus file was imported to SplitsTree v5 for visualization (Huson and Bryant, 2006). We assessed the differentiation between the inferred clusters using the non admixed individual of each cluster and the *ρ* statistic computation implemented in Genodive v3.0 (Meirmans, 2020). The *ρ* statistic is an analogue of *F_ST_* adapted for polyploid data Meirmans et al. (2018). Note that *ρ* values tend to be higher than *F_ST_* values, although both range between zero and one. To unravel the level of genetic diversity, the inferred genetic groups were compared for several diversity estimates using non-admixed individuals. In details, we calculated the observed gametic heterozygosity, the gene diversity and the inbreeding coefficient (G*_is_*) for each genetic group using Genodive v3.0 (Meirmans, 2020; Meirmans et al., 2018). We also utilized ADZE v1.0 to estimate the allelic richness and the private allelic richness through a rarefaction procedure (Szpiech et al., 2008) and their standard deviation. The maximum standardized sample size used for the calculation (i.e. the G ADZE parameter) was set to 12. This corresponded to the number of alleles present at each locus in the smallest groups (i.e. three individuals and four alleles per individual for a given locus).

Genetic structure and diversity were also investigated across different geographical regions because a site-based analysis was impossible given the very low number clonal lineages identified per site. While visual attribution of sites to a region, as done in Bina et al. (2022) was an option, it would not have been entirely reproducible and might have induced inconsistencies in the criteria defining a regional group. To address this, we used a method driven solely by the GPS location of each site, constructing a spatial graph and identifying clusters of geographically close sites using community detection algorithms. In detail, we determined the pairwise geodesic distances between sites using the distm function of the R package geosphere v1.5.18 (Hijmans, 2022).

We then constructed spatial graphs, using the sites as nodes, with the R package igraph v1.4.1 (Csardi and Nepusz, 2006). In order to compare French populations with other *R. gallica* populations, we independently created two graphs, one for France and one for other wild sites, using the function graph_from_adjacency_matrix. The construction of the graphs required setting a threshold above which two nodes are considered not connected. After testing various values, a threshold of 165 km was retained as it effectively distinguished the most expected regions. Sites co-occurring in the same region were then determined by performing graph-based community detection (i.e., identification of spatial clusters) separately on the two graphs. The community detection was performed using the igraph function cluster_infomap with 500 trials and weighted edges (Rosvall and Bergstrom, 2008). Each detected community was named according to its regional origin and hereafter referred to as a regional group. Finally, the community detection sorted the sites into 5 French and 14 European regions (Fig. 3, Table S6). The pairwise *ρ* statistics between regions were calculated using Genodive v3.0 (Meirmans, 2020). Significance was assessed using 10 000 permutations. Regions with fewer than five clonal lineages were excluded from the analysis (i.e., Spain, Moldova, North Czechia, West Romania). A second community detection analysis was performed on a graph built from the pairwise *ρ* statistics between regions. In this graph, regional groups were represented as nodes, and edges connected pairs of regions showing low differentiation (*ρ <* 0.05). The analysis, conducted with the same igraph (Csardi and Nepusz, 2006) cluster_infomap settings as described above, aimed to identify clusters of regions that were genetically similar based on low pairwise differentiation. Additionally, using ADZE, we calculated the private allelic richness per region and tts standard deviation. The G ADZE parameter was set to 16, and regions containing less than four clonal lineages were excluded from the analysis (i.e. Spain, Moldova, North Czechia).

**Figure 3.**
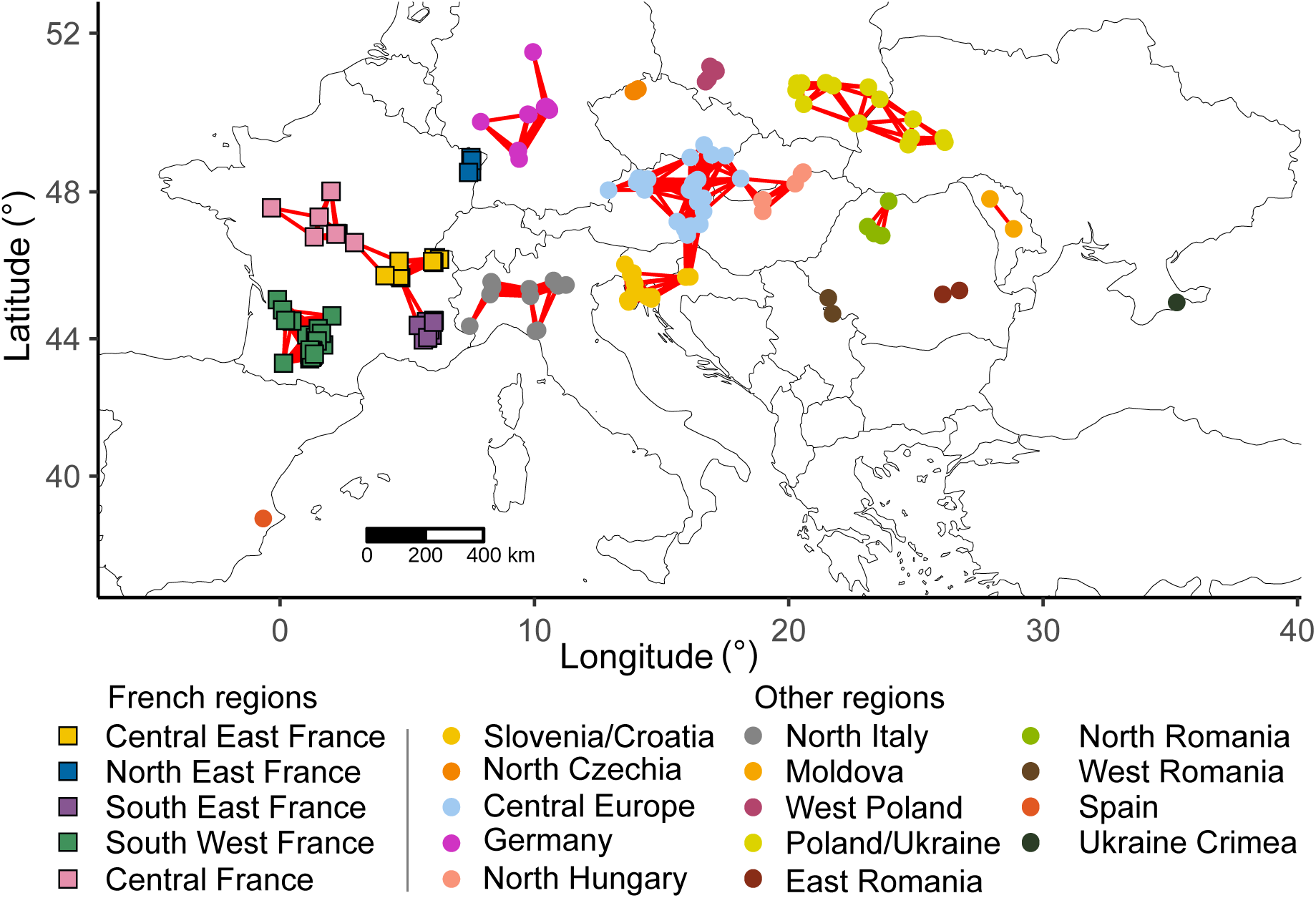
Spatial networks of collection sites for the wild individuals present in the *R. gallica* dataset. Nodes of the network computed for French sites are represented as squares. Nodes of the network computed with for non French sites are represented by circles. Colors indicate regional groups inferred by community detection. Red edges indicate a geographic distance between two sites below the threshold of 165 kilometers.

## Results

### Ploidy and marker quality check

After the removal of the four suspected paralogs, filtering for error rates and optimizing for missing data, the *Rosa* spp. dataset was composed of 29 loci and 1618 samples. Each of the seven chromosomes was covered by at least three loci (Fig. S5). Wild-collected *R. gallica* in this dataset represented 219 out of the 223 collection sites, meaning that we failed to genotype individuals from four sites (Fig. 1). The four missing sites were all located in France and were represented each by only one herbarium specimen. Our ploidy prediction model achieved an accuracy of 89.6% according to the out-of-bag estimates. Of the 1618 individuals, 140 were considered diploid, 1381 were classified as tetraploid, and 97 as hexaploid. Ploidy estimation results are available in Table S7. The average error and missing rates of the 29 loci were 2.17% and 5.42% respectively (Table 1). A total of 2326 and 884 alleles were identified in the *Rosa* spp. (Table 1) and *R. gallica* (Table S8) datasets respectively. Length homoplasy was quite common, as the number of alleles increased by 193.3% when considering whole amplicon information compared to solely considering allele length for the *Rosa* spp. dataset. For the *R. gallica* dataset, a more moderate increase of 80% was observed. Length homoplasy occurred even for markers for which alleles were called by focusing on the repeat motif only (Table 1). On the *Rosa* spp. dataset, the PIC was high, ranging from 0.59 to 0.95 (Table 1), highlighting the good discriminatory power of the markers. The investigation for null alleles revealed a significant correlation between F*_is_* and F*_st_*, a correlation between F*_is_* and missing rates, and the standard deviation of F*_is_* is four times that of F*_st_* (Appendix A). This seems to indicate that null alleles may affect the data. However, as most of the loci showed an excess of homozygotes, removing the ones with this excess would have removed a large part of the loci. Additionally, such a high number of loci with an excess of homozygotes suggested that other factors than only the null alleles may be also responsible for the excess. Therefore, we decided to keep the 29 loci and to take caution during the interpretation of differentiation values as these can be inflated by the presence of null alleles.

**Table 1.** Quality and polymorphism of the 29 loci, assessed on the *Rosa* spp. dataset. N_WAI: Number of alleles called by considering the Whole Amplicon Information. N_length: Number of alleles considering only allele length. Rare allele: Percentage of alleles, detected by considering the Whole Amplicon information, with a frequency lower than 1%. PIC: Polymorphism Information Content.

| Locus | Calling strategy | Missing rate (%) | Error rate (%) | N_WAI | N_length | Rare allele (%) | PIC |
| --- | --- | --- | --- | --- | --- | --- | --- |
| Gal01_03 | RepeatFocused | 10.14 | 2.84 | 80 | 33 | 81.3 | 0.61 |
| Gal02_11 | FullLength | 10.07 | 3.10 | 109 | 46 | 79.8 | 0.92 |
| Gal02_14 | FullLength | 5.75 | 2.21 | 233 | 54 | 83.3 | 0.95 |
| Gal03_19 | FullLength | 2.41 | 0.94 | 50 | 27 | 56.0 | 0.86 |
| Gal03_25 | RepeatFocused | 1.79 | 1.40 | 45 | 17 | 80.0 | 0.60 |
| Gal04_34 | RepeatFocused | 0.74 | 2.42 | 50 | 24 | 68.0 | 0.80 |
| Gal05_38 | FullLength | 14.52 | 2.84 | 96 | 24 | 79.2 | 0.70 |
| Gal06_42 | RepeatFocused | 15.45 | 2.17 | 86 | 33 | 65.1 | 0.91 |
| Gal07_48 | RepeatFocused | 4.88 | 1.69 | 49 | 24 | 67.3 | 0.85 |
| Gal07_51 | RepeatFocused | 0.93 | 0.51 | 44 | 18 | 75.0 | 0.73 |
| Gal07_55 | FullLength | 8.47 | 2.10 | 97 | 28 | 77.3 | 0.77 |
| Gal07_56 | FullLength | 9.02 | 2.05 | 97 | 24 | 68.0 | 0.89 |
| Lab01_H9B07_07 | RepeatFocused | 0.19 | 2.16 | 31 | 15 | 71.0 | 0.79 |
| Lab03_Rh58_24 | FullLength | 6.12 | 0.84 | 88 | 20 | 89.8 | 0.62 |
| Lab07_Rw5G14_54 | RepeatFocused | 0.74 | 2.17 | 66 | 24 | 68.2 | 0.89 |
| Trf01_04 | RepeatFocused | 0.06 | 1.34 | 45 | 18 | 64.4 | 0.82 |
| Trf01_05 | RepeatFocused | 0.12 | 0.61 | 28 | 15 | 67.9 | 0.59 |
| Trf02_08 | RepeatFocused | 0.68 | 1.68 | 60 | 29 | 56.7 | 0.90 |
| Trf03_21 | RepeatFocused | 0.37 | 1.13 | 93 | 25 | 77.4 | 0.86 |
| Trf03_23 | FullLength | 0.87 | 1.37 | 62 | 33 | 74.2 | 0.64 |
| Trf04_27 | FullLength | 14.52 | 3.67 | 108 | 38 | 75.9 | 0.93 |
| Trf04_30 | RepeatFocused | 11.37 | 3.26 | 52 | 17 | 59.6 | 0.86 |
| Trf05_35 | FullLength | 11.37 | 2.78 | 116 | 31 | 81.9 | 0.85 |
| Trf05_39 | RepeatFocused | 1.55 | 2.76 | 36 | 22 | 44.4 | 0.75 |
| Trf06_44 | RepeatFocused | 9.46 | 4.46 | 124 | 39 | 68.5 | 0.95 |
| Trf06_47 | FullLength | 0.62 | 1.49 | 107 | 24 | 73.8 | 0.84 |
| Trf07_50 | RepeatFocused | 8.22 | 4.60 | 41 | 28 | 63.4 | 0.83 |
| Trf07_52 | RepeatFocused | 4.82 | 3.65 | 147 | 40 | 71.4 | 0.95 |
| Trf07_53 | RepeatFocused | 1.85 | 0.74 | 86 | 23 | 60.5 | 0.91 |
| Average |  | 5.42 | 2.17 | 80.21 | 27.34 | 70.67 | 0.81 |

### Wild-cultivated clonal lineages

The distance-based method used for clonal lineage detection identified clonal relationships for 1041 out of the 1618 successfully genotyped samples. After retaining a single representative per clonal lineage and including non-clonal individuals, a total of 886 distinct clonal lineages were identified. These lineages ranged from single-individual (non-clonal) to multi-individual (clonal) lineage. Additionally, the method has a good specificity as no clonal relationship was found among the 9 half and 10 full-sib successfully genotyped individuals. After removing these 19 sibs, the clone-corrected *Rosa* spp. dataset was composed of 867 individuals (Table S2). We detected clonal lineages within the same site for 163 out of the 195 sites for which we successfully genotyped more than one individual. Moreover, 36 out of the 219 sites contained at least one individual that belonged to the same clonal lineage as either a cultivated (or botanical) individual or a wild individual collected from a different site (Fig. 4 and Table 2). Two clonal lineages included only individuals from different wild sites (Fig. 4). One contained all individuals from three Spanish sites (clonal lineage J) and the other contained three individuals from three Slovenian sites (clonal lineage K) that were sampled in the botanical garden of Ljubljana. Two samples collected in Bosnia and Herzegovina were clonal with cultivated individuals (clonal lineages C and F). In France, clonal lineages regrouping botanical or cultivated individuals and wild sampled individuals were detected across 28 sites out of the 98 French sites (clonal lineages A, B, D, E, G, H and I). The largest and most widespread clonal lineage (B, Table 2) contained individuals distributed at 13 sites in Central and Western France (Fig. 4). Individuals of the clonal lineages A and G were collected from six and five sites, respectively. Other lineages were less extended as they contained individuals originated from three (clonal lineage I), two (clonal lineage D) or one site (clonal lineages E and H, Fig. 4). The largest clonal lineage (B), included one botanical individual called *R. gallica* var. *versicolor* and two cultivars named ‘Rosiers des parfumeurs’ and ‘Bouquet de Vénus’. The flower phenotypes of individuals belonging to the clonal lineage B showed a number of petals well above the petal number usually found in sites where no clonal relationship was found with cultivars or with specimens of other wild sites (Fig. 5).

**Figure 4.**
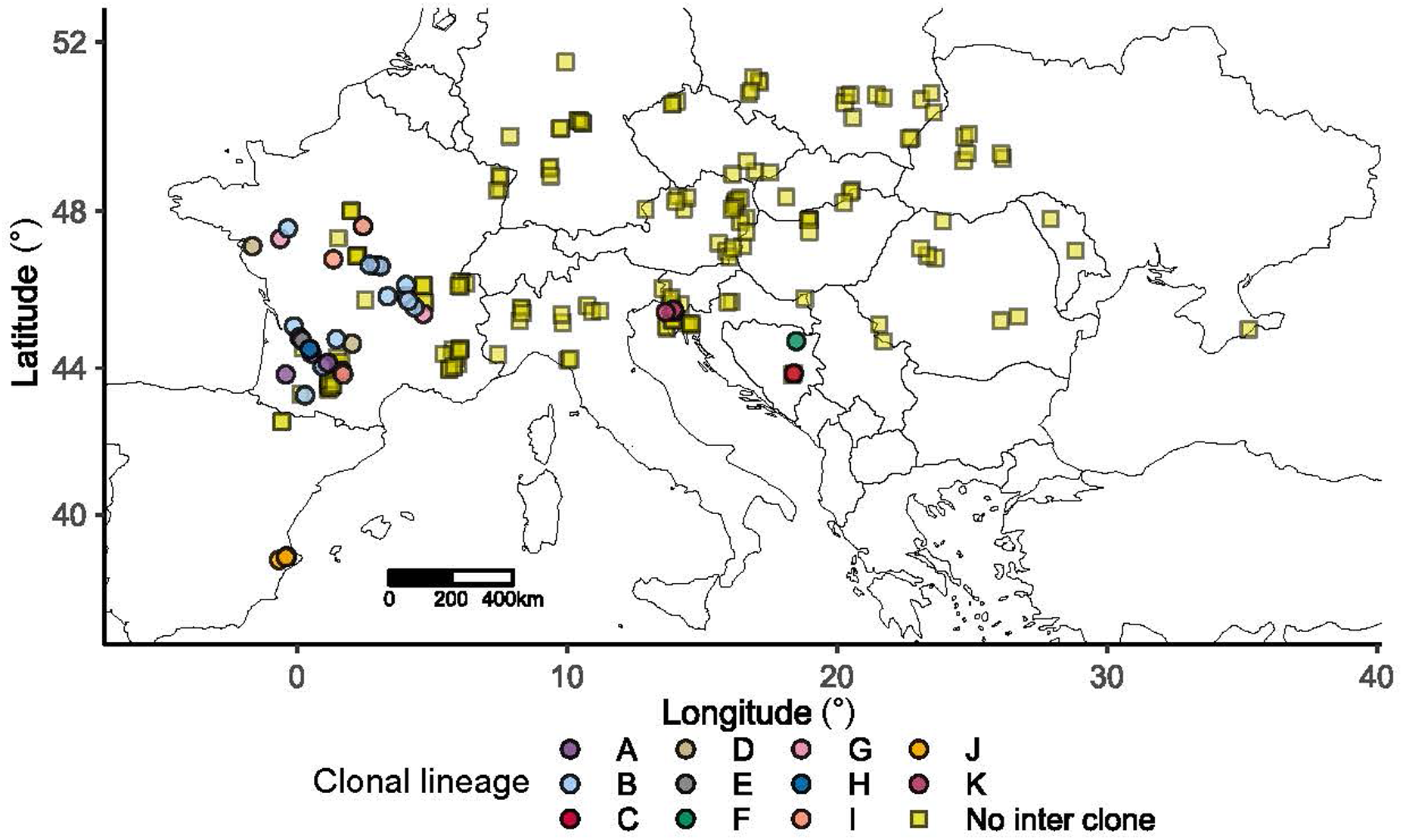
Clonal relationships between individuals from different sites and/or between wild and botanical or cultivated individuals. The colors indicate the detected clonal lineage. Only wild sites are presented on the map. A site is colored as belonging to clonal lineages A to K, if at least one individual from that site belongs to one of these clonal lineages. Botanical or cultivated individuals belonging to the different clonal lineages are: (A) ‘Agathe Rose’ - ‘Rose de Rescht’, (B) *R. gallica* var. *versicolor* - ‘Bouquet de Vénus’ - ‘Rosier des parfumeurs’, (C) ‘Frankfurt’ - *R. gallica* var. *splendens*, (D) ‘La belle Sultane’ - *R. centifolia* ‘Simplex’ - ‘Rosa violacea’, (E) ‘Parure des vierges’, (F) ‘Roi des Pays-Bas’ - *R. gallica* ‘Agatha’ - ‘Rosier d’amour’, (G) *R. centifolia* ‘Parvifolia’ - *R. centifolia* ‘Pomponia’, (H) Unnamed cultivars, (I) a botanical *R. gallica*. The clonal lineages J and K contain individuals from three Spanish or three Slovenian sites, respectively. “No inter clone” indicates that, for the site, no clonal relationship was detected with botanical or cultivated individuals or with individuals for other wild sites.

**Figure 5.**
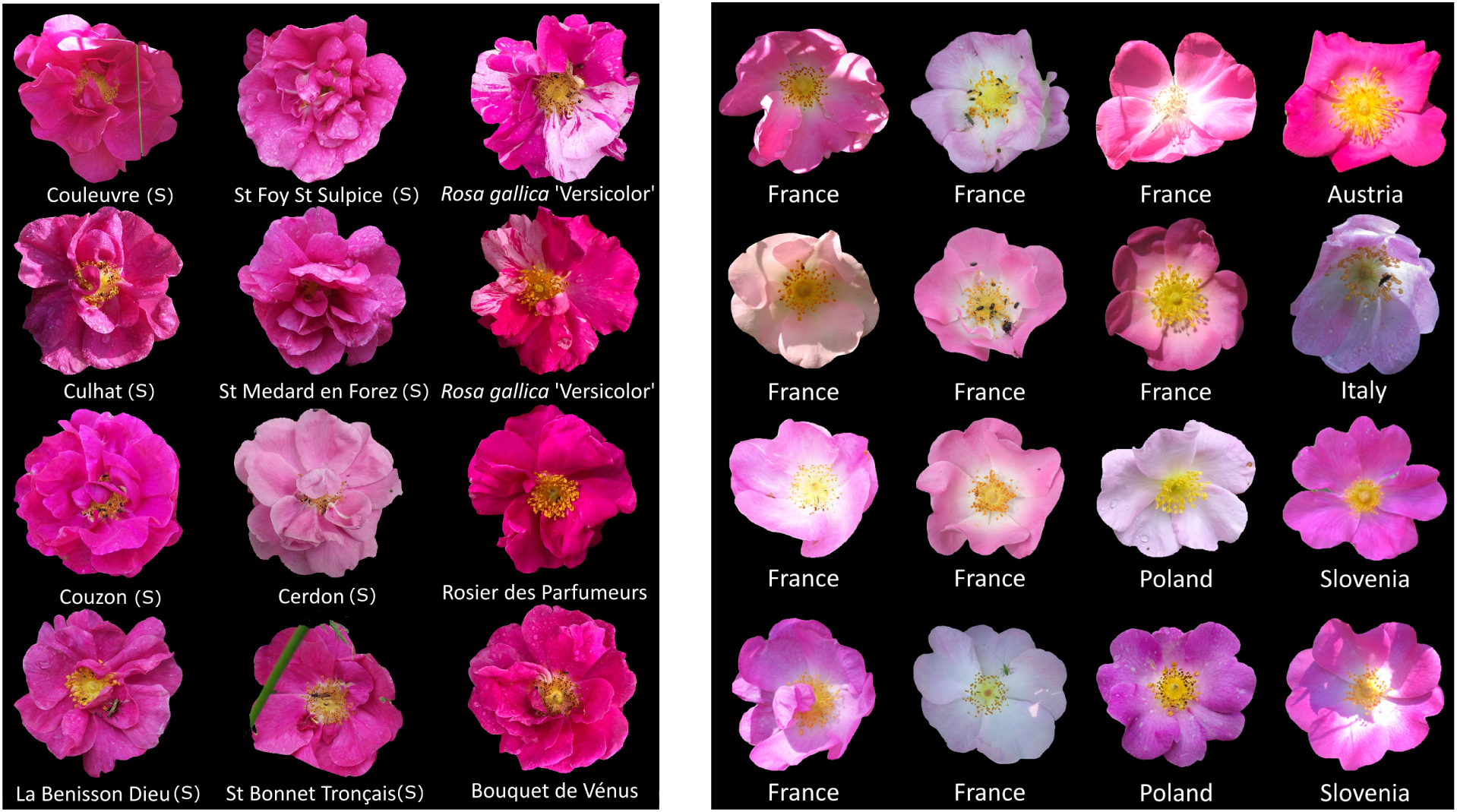
Comparisons between flowers of 12 individuals belonging to clonal lineage B (left) and flowers found in sites for which no individual showed any clonal relationship with cultivars or individuals from other sites (right). The name of the cultivars or of the sampling site (S, left part) or of the country of the site (right part) are indicated below each flower picture. Credits: Genetic and Diversity of Ornamentals (GDO) research team, Institute of Research in Horticulture and Seeds, France, Clovis Pawula, Alexander Mrkvicka, Jordane Cordier, Renata Piwowarczyk and Fabrizio Bonali.

**Table 2.** Number of samples per clonal lineage.

| Clonal lineage | Number of samples | Clonal lineage | Number of samples |
| --- | --- | --- | --- |
| A | 22 | F | 4 |
| B | 57 | G | 24 |
| C | 3 | H | 2 |
| D | 15 | I | 9 |
| E | 3 | J | 9 |
| K | 3 |  |  |

### Mis-identification detection

PCoA analyses (Appendix B.A1–A2) showed grouping by botanical section. Indeed, most *R.gallica* grouped together, although some individuals clustered with *Caninae* or European *Synstylae*, consistent with PCA results (Appendix B.B1–B2). Subsenquently, a DAPC was used to statistically determine which individuals belonged to the *R. gallica* species. The probability of being a *R. gallica*, determined based on the first two discriminant functions at *K* = 12 (Appendix C) and on the DAPC genetic groups content (Appendix D), revealed 27 individuals, sampled as being *R. gallica* specimens, that are actually not belonging to this species or are hybrids (Appendix E). Among these, 19 were grouped with *Caninae*, two with European *Synstylae*, and one with *SynstylaeIndicae*. Three others were distributed among other non-*R. gallica* groups that regrouped individuals from different botanical sections or horticultural groups. The remaining two were part of *R. gallica* groups but with membership probabilities below the threshold suggesting a hybrid origin (Appendix E and Table S9). In total, 519 individuals were identified as belonging to the *R. gallica* species. These included 415 wild-collected *R. gallica*, 97 botanical or cultivated presumably related to *R. gallica* and 7 individuals that were initially collected as not being related to *R. gallica* (these were considered as being either botanical or cultivated for further analyses depending on historical information available, Table S1, S2). These 519 individuals constituted the *R. gallica* dataset and were used to investigate the European diversity of the species and the origins of French populations.

### *R. gallica* structure and diversity

In both multivariate analyses (i.e. PCoA on k-mer dissimilarities and PCA on intra-individual allele frequencies), the wild individuals from other countries than France (red circles) were grouped on the right of the plots while botanical or cultivated (blue triangles) and individuals from South Western France (green diamonds) were on the left of the plots (Fig. 6). The second principal component of the PCoA (Fig. 6.A1) and the third principal component of the PCA (Fig. 6.B2) did not separate clearly the individuals according to their nature or sampling site location. On the PCoA plot showing the first and third principal component (Fig. 6.A2), the South Western France individuals (top left, green diamonds) tended to be separated from the non French individuals (top right, red circles) and from the botanical or cultivated specimens (blue triangles). The same clear separation was observable on the first two principal components of the PCA (Fig. 6.B1) and the three groups seem even more separated. When looking at the same plot (Fig. 6.B1), individuals from other French regions (yellow diamonds) were either grouped with the specimens sampled in South Western France or with the ones sampled in non French sites. More specifically, individuals from the East of France (North and South regions) were mostly grouped with wild-sampled specimens from other countries (Appendix F). Individuals from Central France were mostly found in intermediate locations between the group from South Western France and the group of individuals sampled in other countries (Appendix F).

**Figure 6.**
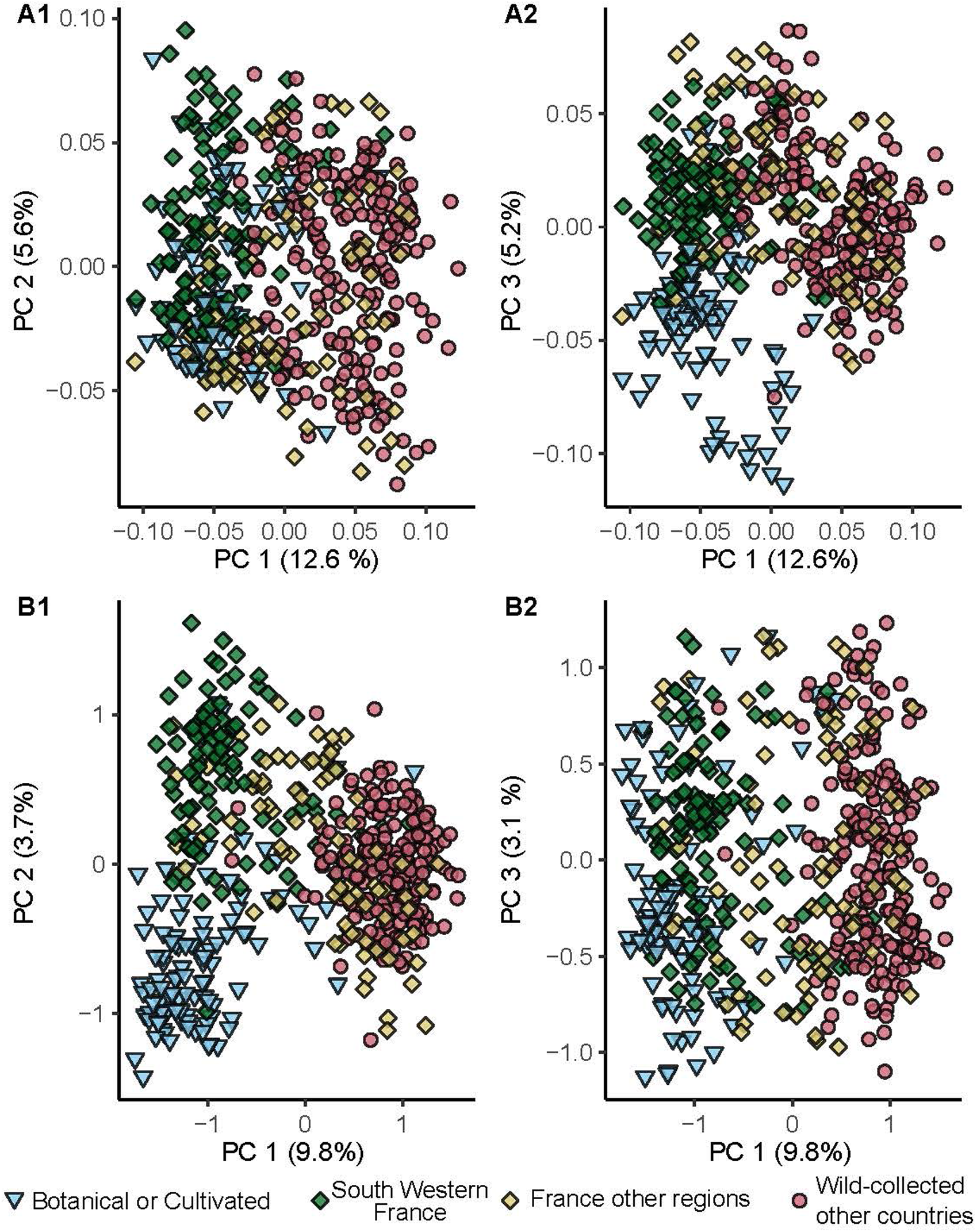
Multivariate analyses of the *R. gallica* dataset. (A) PCoA based on pairwise k-mer dissimilarities. (B) PCA based on intra-individual allele frequencies. A1 and B1 correspond to individual coordinates for the first and the second principal components while A2 and B2 show the first and the third principal components whatever the multivariate analysis. Cultivated or botanical individuals are depicted as triangles, whereas French wild-collected ones are shown as diamonds, and other wild-collected individuals are represented by circles.

In order to unravel the diversity structure, quantify differentiation, and identify admixed *R. gallica* individuals, the inferential model-based software STRUCTURE was used. The ΔK and the parsimony-based methods, respectively, suggested K = 2 and K = 8 as probable optimal numbers of clusters, although K = 7 had also high parsimony index value (Fig. S6). The major mode was well supported at K = 2 (Appendix G) and K = 8 (Fig. 7). At K = 2, almost all cultivated individuals, half of the botanical ones, and wild individuals from South Western and Central France were grouped together, along with the the clonal lineage from Spain (Appendix G). Additionally, some individuals from Central Eastern France appeared admixed between the two groups. *Rosa gallica* individuals from other countries were grouped with individuals from the East of France and the remaining half of botanical ones (Appendix G). At K = 8, France appeared highly structured in comparison to the rest of the range. Indeed, while individuals from the rest of the range were almost exclusively assigned to only two clusters (grey and orange clusters, Fig. 7), French individuals were distributed among seven genetic clusters, five of which were almost exclusively found in France. Moreover, the K = 8 inference revealed a spatial genetic structure (Fig. 7). Indeed, all inferred clusters regrouped individuals from one or several adjacent regions. The widest cluster (orange), located in Central Eastern Europe, covered almost half of the area studied. In detail, individuals belonging to this cluster occurred from north east of France to Moldova and from Poland to the north of the Balkans. Individuals from South Alps (i.e. North Italy, Slovenia/Croatia, and South East of France) mainly belonged to the grey cluster or were admixed between the orange and the grey clusters. Another cluster (light blue) regrouped almost exclusively individuals from South Western and Central France, with one exception from Spain. Two well-defined clusters were formed by individuals from Central France (pink) and individuals from Crimea (dark orange, Fig. 7A-B). A few individuals from South Western France (dark blue) and Central Eastern France (green) formed two restricted clusters. Almost all cultivated individuals and half of the botanical ones belonged to the same cluster (yellow, Fig. 7A-B). Individuals from three wild sites, two from South Western France and one from Central Eastern France, were part of this cultivated cluster. In the STRUCTURE analysis, clonal lineages including individuals from different sites and cultivated or botanical ones were represented by one individual. These ones mostly belonged either to the cultivated cluster (yellow) or to the cluster regrouping individuals from South Western and Central France (light blue, Fig. 7A).

**Figure 7.**
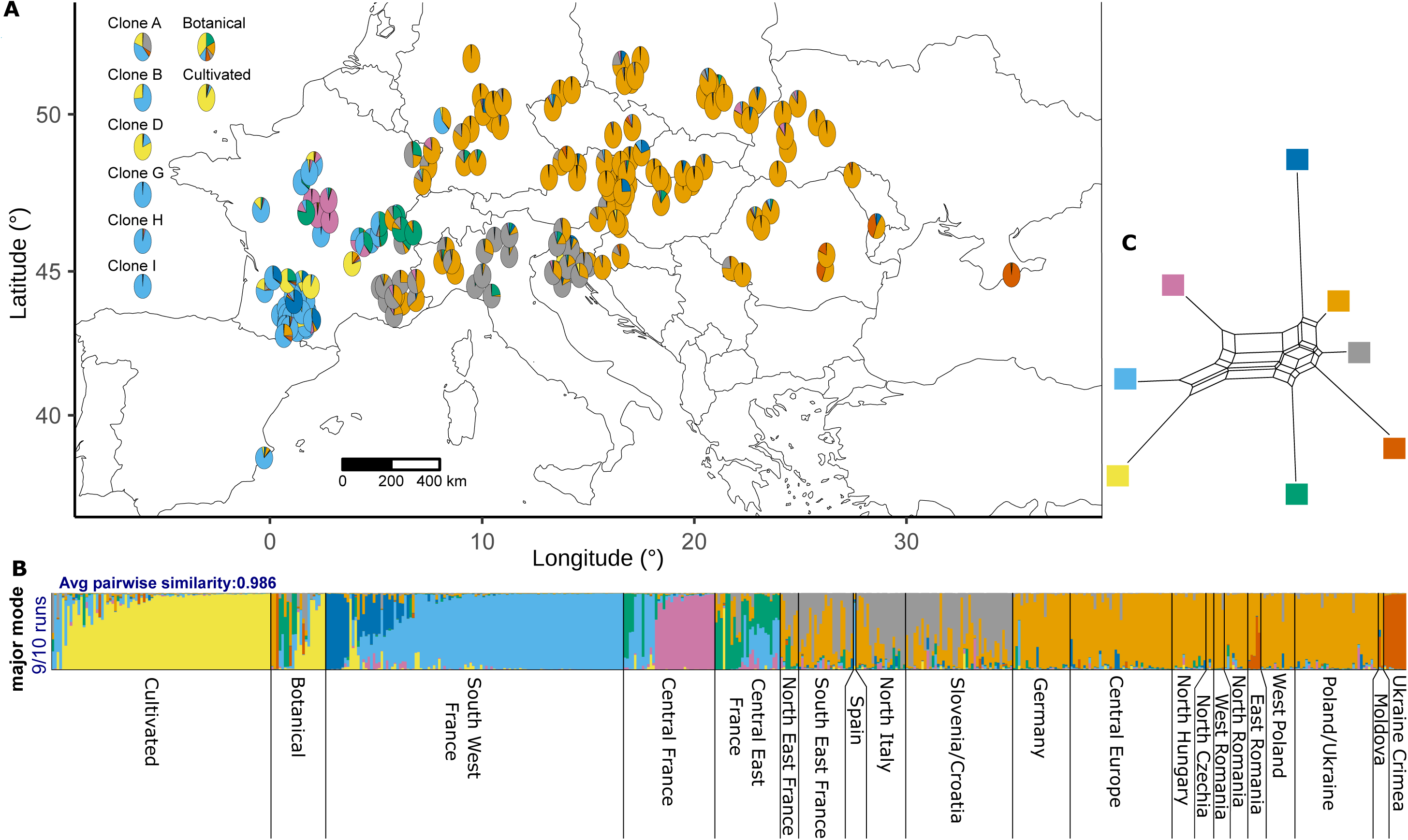
Inferred population genetic structure of *R. gallica* at K = 8 (N = 519). (A) Mapping of mean ancestry proportions per site of collection. Ancestry proportions of the representative individual of the clonal lineages A, B, D, G, H and I, are shown on the left of the map. Mean ancestry proportions of cultivars and botanicals are also available. The site coordinates were slightly jittered to prevent overlapping circular diagrams and improve map readability, the exact mapping is available in figure S7. (B) STRUCTURE barplot representing the major mode found using Pong v1.5 (Behr et al., 2016). (C) Unrooted split network showing allele frequency divergence among STRUCTURE clusters.

The split network, visualizing differentiation through allele frequency divergences among inferred genetic clusters, displayed two primary partitions. The first comprised clusters of cultivated specimens, South Western France, and Central France wild individuals (represented by yellow, light blue, and pink respectively). The second partition exclusively included clusters of wild sampled plants (dark orange, orange, grey, green, and dark blue, Fig. 7C). Notably, the most extensive cluster found in South Western and Central France (light blue) was genetically closer to the cultivated cluster (yellow) than to any other clusters. The cluster primarily found in Central France (pink) was closest to the clusters from the second partition among all clusters from the first. The clusters regrouping individuals from Central Eastern Europe (orange) and from South Alps (grey) showed low divergence (Fig. 7C). The geographically limited cluster from South Western France (dark blue) showed relatively high divergence, similar to the clusters from Central Eastern France and Crimea (green and dark orange respectively). The *ρ* statistics revealed that all clusters were significantly differentiated from each other (p-values < 0.05, 10 000 permutations, Fig. 8D). Values of *ρ* varied from 0.067 (clusters from Central Eastern Europe, orange, and from South Alps, grey) to 0.491 (clusters from Central Eastern France, green, and from South Western France, dark blue). The green cluster showed high degree of differentiation with almost all other clusters while the grey and orange cluster were the lowest differentiated on average (Fig. 8D). Globally, *ρ* values were in accordance with the split networks partitions (Fig. 7 and Fig. 8D). When looking at the *ρ* differentiation between geographical regions, the community detection revealed eight communities on the differentiation graph (Fig. 9 and Table S10). One community spanned from South East France to Croatia/Slovenia and corresponded well with the STRUCTURE cluster of South Alps (grey, Fig. 9). Another community regrouped regions from Central Germany to North Romania and therefore corresponded well to the STRUCTURE cluster from Central Eastern Europe (orange, Fig. 9). The six others corresponded to isolated regions.

**Figure 8.**
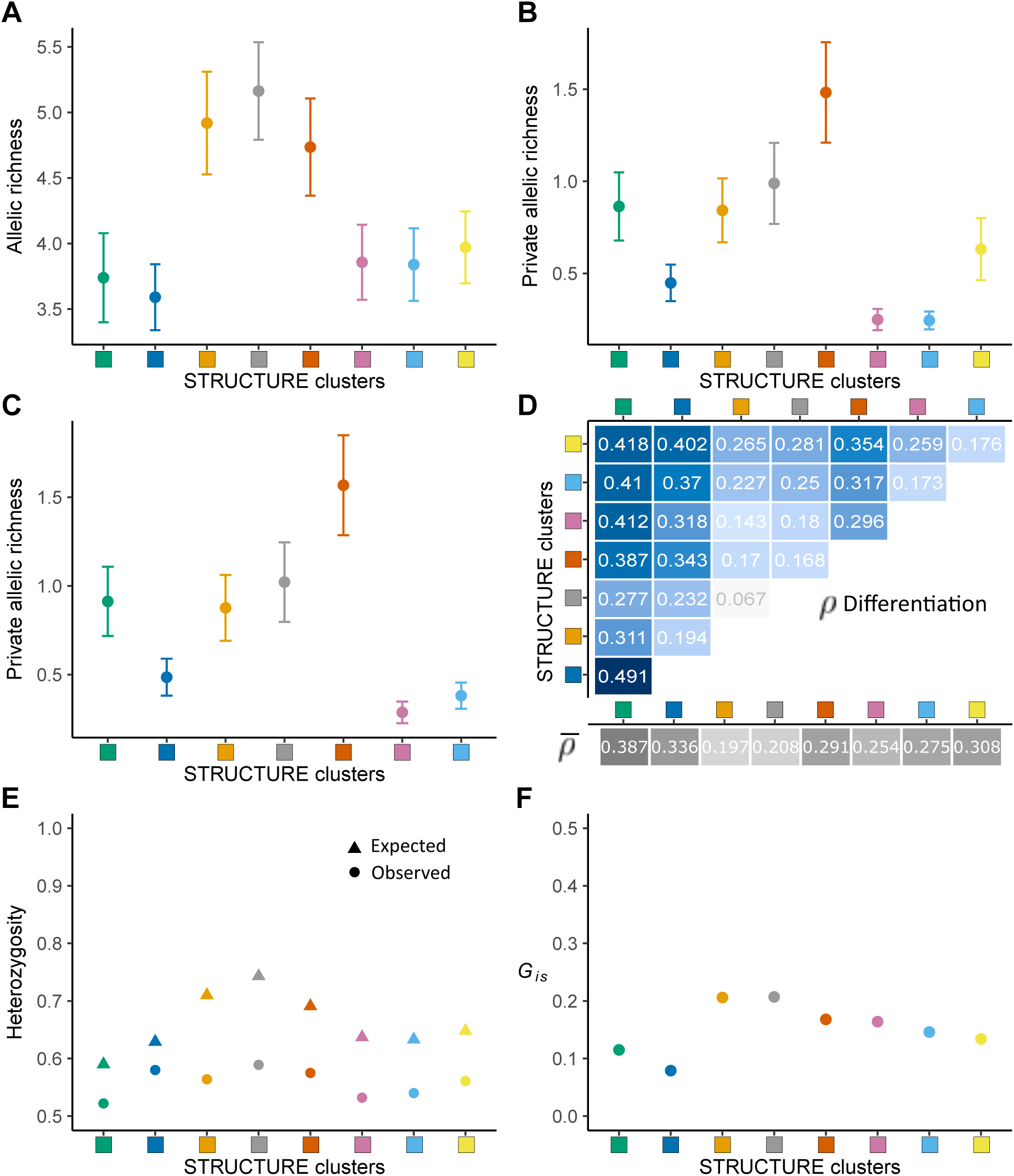
Diversity indices and differentiation among STRUCTURE clusters at K = 8. (A) Sample size independent mean allelic richness per locus. (B) Sample size independent mean private allelic richness per locus. (C) Same as B but determined without the yellow cluster. Error bars reflect standard error of the mean across loci. (D) Pairwise *ρ* differentiation values. Blue intensity is proportional to the *ρ* value. Average *ρ* are indicated in grey cells (E) Expected and observed heterozygosity. (F) Inbreeding coefficient (G*_is_*).

**Figure 9.**
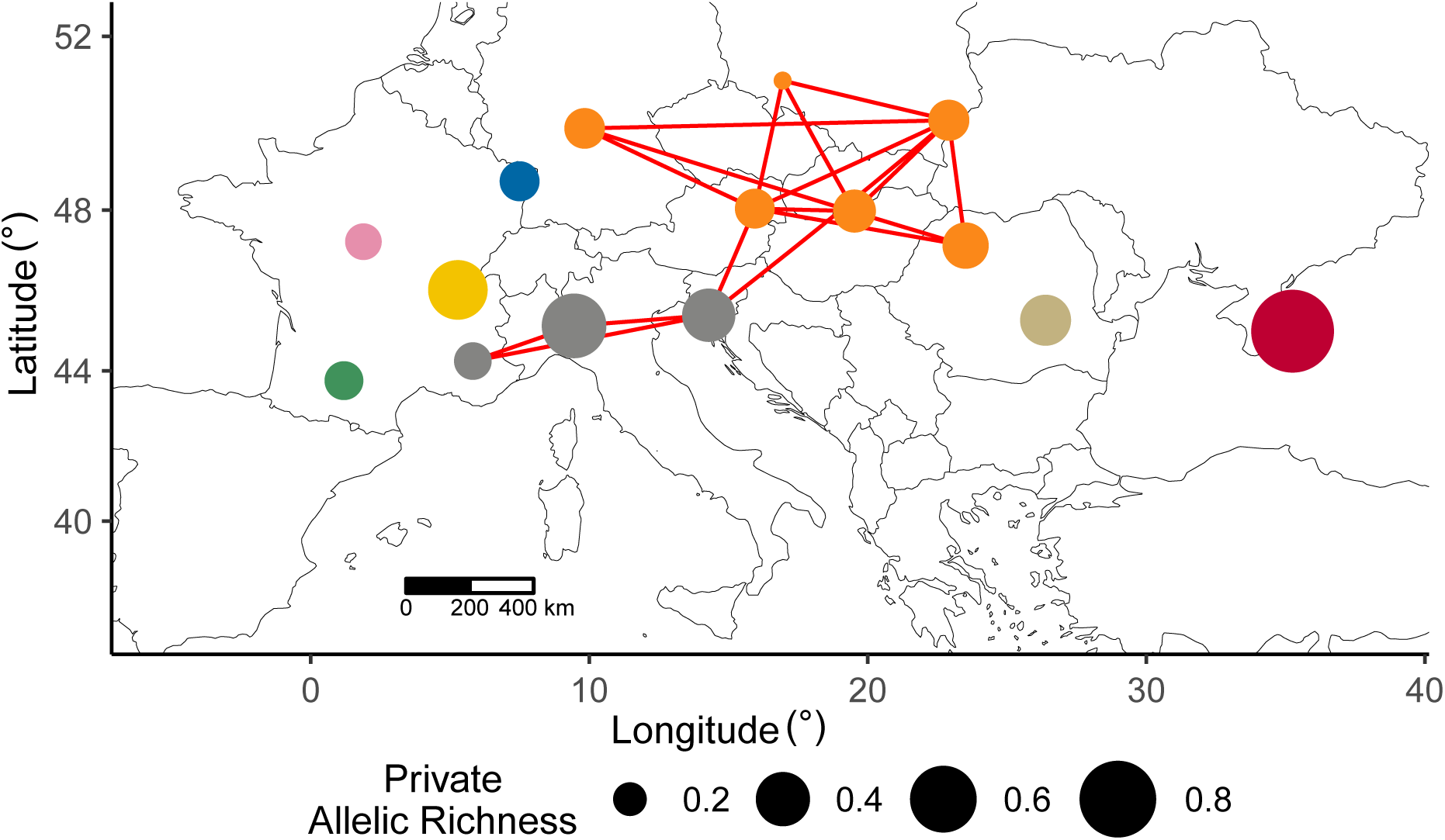
Graph of pairwise *ρ* differentiation and private allelic richness across geographical regions. Edges connect regions with a *ρ* value lower than 0.05, while nodes represent geographical regions. Node colors indicate detected groups of weakly differentiated regions (graph-based communities). Node size is proportional to private allelic richness, estimated with ADZE using a G parameter of 16. Regions represented by fewer than four clonal lineages were excluded from the analysis.

Clusters of individuals from South Alps, Central Eastern and Eastern Europe (grey, orange, and dark orange) had higher allelic richness than other clusters (Fig. 8A). The cluster from Crimea (dark orange) reached the highest private allelic richness. Very low private allelic richness was observed for groups growing almost exclusively in France, namely the dark blue, pink, and light blue clusters (Fig. 8B). The private allelic richness was also calculated without the cultivated cluster (yellow) to focus on wild-collected individuals. Even in this case, the private allelic richness of the the dark blue, pink, and light blue clusters remained low (Fig. 8C). When looking at the diversity per geographical region, the highest private allelic richness was recorded in Crimea (Fig. 9 and Table S11). Other high private allelic richness values were found for Central Eastern France and North Italy. On the contrary, the private allelic richness levels in Central and South Western France were among the lowest (Fig. 9). The expected heterozygosity was higher in clusters of individuals from South Alps, Central Eastern and Eastern Europe (Fig. 8E). Inbreeding coefficient (*G_is_*) values exhibited minor variations among groups. However, *G_is_*was the lowest in the small cluster of individuals from South Western France (dark blue). The highest *G_is_* values were observed in the two largest groups (orange and grey, Fig. 8F).

## Discussion

In the present study, we investigated the range-wide genetic diversity and structure of *R. gallica* with a particular focus on French wild populations. We genotyped a large panel of *R. gallica* and other *Rosa* spp. individuals using a recently developed SSRseq marker set (Pawula et al., 2023). We defined an analysis framework adapted to analyze diversity and structure of a species with both axesual and sexual reproduction, being part of a genus with high rate of heterozygosity, interspecific hybridization and polyploidization. Indeed, we sequentially selected high quality loci, retained one individual per clonal lineage and removed mis-identifications to focus on *R. gallica*. Then, analyzing the largest collection of specimens of this species, we revealed the presence of a spatial genetic structure over the species range. The greater allelic diversity found in South Alps and Central Eastern Europe compared with the one found in the most Western part of the species range and the higher private allelic richness found in low latitude locations may suggest that a natural recolonization took place since the last glacial maximum. Moreover, the strong genetic structure found in France and the low allelic diversity found in Western France could be explained by a more recent colonization of this country part mediated by natural and human factors. The multiple clonal relationships found between cultivars and wild-collected *R. gallica* are clear evidences of recent human-mediated dispersal, although this is only certain for the sites where we found such clonality.

### Highly structured French populations

The French wild populations appear more structured than those from the rest of the range, as evidenced by the higher number of genetic clusters detected within France and the strong differentiation observed among the five French geographical regions considered. Individuals from South Western, Central, and Central Eastern France were distinct from those in Northern and Southeastern France, the latter grouping with individuals from the South Alps and Central Eastern Europe within the same two clusters. This pattern suggests that some French populations share a common demographic history with foreign populations, while other French populations (from South Western, Central, and Central Eastern regions) might have undergone different or additional demographic events. Notably, the clusters in these areas exhibit low allelic richness and low private allelic richness, indicating that these populations might share the same origin as those from South Alps and Central Europe, but that subsequent migration events could have triggered a genetic bottleneck. Based on these observations, we will examine hypotheses proposed in literature in light of our results, separately for South Alps and Central Eastern Europe (including South Eastern and North Eastern France) and for the rest of the French regions (including Spain which is only represented by a single clonal lineage). As a reminder, these hypotheses include a natural recolonization from refugia after the last glacial maximum, or contributions of human-mediated dispersal at different periods (Antiquity, Middle Age or more recently).

### R. gallica present days structure pattern suggests recolonization from primary refugia

The natural recolonization from Southern refugia (i.e., the Balkans, Iberia, and/or Italy) has been described for several species (Daneck et al., 2016; Magri et al., 2006). Recolonization can occur from one or multiple refugia, leading to different patterns of present-day diversity (Ali, 2020). For instance, when refugia contribute unevenly to the recolonization, a large part of the recolonized area is covered by related populations that appear to belong to the same genetic cluster or to minimally differentiated ones. This phenomenon has been described for *Fagus sylvatica* (Magri et al., 2006) and the meadow grasshopper, *Chorthippus parallelus* (Korkmaz et al., 2014). Additionally, the recolonization process may involve the loss of alleles in recolonized areas. The level of differentiation and the intensity of allele loss depend on the number of founders and the extent of gene flow between the refugium and the recolonized area (Ali, 2020). Here, two clusters cover most of the South Alps and Central Eastern Europe (i.e., clusters orange and grey) and this predominance of two clusters was confirmed by the community detection performed on the *ρ* differentiation graph. Moreover, low latitude regions of this area seem to exhibit more private allelic richness than high latitude regions. This is consistent with a loss of diversity between refugia in the south and recolonized areas in the north. Therefore, the present-day diversity pattern of *R. gallica* seems compatible with the hypothesis of a recolonization following the last glacial maximum. Determining from where the South Alps and Central Eastern Europe were recolonized remains highly speculative. Here, the two clusters are only minimally differentiated and low differentiation is present between the geographical regions they cover. Therefore, South Alps and Central Eastern Europe could have been recolonized from one unique refugium (as described for *Fagus sylvatica*, Magri et al., 2006). As for these regions the highest private allelic richness is found in North Italy, the refugium could have been the Italian peninsula. In this scenario, the Alps could have acted as a geographical barrier responsible for the minimal but significant differentiation between the grey and orange clusters. This would also explain the low differentiation found between Croatia/Slovenia and two regions from Central Europe. Cryptic northern refugia have been suggested for another European *Rosa* species, *R. pendulina* (Daneck et al., 2016). However, we did not detect any indication of a northern refugium for *R. gallica*, as no high-latitude region exhibited high private allelic richness. Finally, the third cluster found in Eastern Europe (the dark orange one) is located at the extreme Eastern part of the studied range. This cluster shows high diversity and private diversity, suggesting that another refugium might be present in this area, but without a significant westward post-glacial expansion.

The other hypotheses mentioned humans as important factors in the expansion process of *R. gallica*, either during the Romans age (Debray, 2020) or during the Middle Age (Saakov, 1965). The Romans cultivated few types of roses that might eventually be attributed to *R. gallica*. These would have been the rose of Praeneste, the rose of Milet and the rose of Trachyne (Joret, 1892). At this age, the roses were mainly multiplicated by the root suckers (Mattock, 2017). During the Middle age, the propagation by root suckers was still practiced and there was only few varieties of *R. gallica* (Joret, 1892). The clonal propagation of *R. gallica* from a reduced number of distinct individuals by root suckers would have let trace on the actual genetic diversity. Under these human-mediated dispersal hypotheses, few parental genotypes of roses would have been propagated by humans across Europe by clonal reproduction. As hundreds years spent, the clonal relationships might be difficult to retrieve due to potential sexual reproduction generations since that time. However, a clonal propagation at a particular time would have represented a strong bottleneck leading to a low degree of diversity observed within some genetic clusters. Additionally, the expected clustering pattern would be influenced by the number of varieties propagated per region and the potential hybridization between varieties happening after their human mediated dispersal. This could lead to a low spatial genetic structure. Here, we found both a high diversity in the largest clusters (South Alps grey and Central Eastern Europe orange) and a clear spatial genetic structure with low differentiation values found predominantly between adjacent regions. Therefore, our results do not suggest that humans, either during the Romans age or during the Middle age, were important factor of dispersion for *R. gallica* in South Alps and Central Eastern Europe.

### A more recent colonization of South Western and Central France

Only one of the five clusters found in South Western, Central and Central Eastern France is widespread (i.e., the light blue cluster). This one is located in the most Western part of the species range. For several European species, an Iberian or Pyrenean refugium has been described (Daneck et al., 2011, 2016; Magri et al., 2006). Therefore, it would be possible that the Western part of the *R. gallica* range was recolonized from such a refugium. Recolonization from an unsampled Iberian refugium would result in individuals from Western and Central France and Spain belonging to a different genetic cluster than those present in other parts of the range. Furthermore, this cluster would be highly differentiated from others, as the Iberian refugium would have been isolated during the last glacial period. Long-term isolation caused by the prolonged glacial period usually leads to the appearance or loss of different alleles in separated populations (Daneck et al., 2016). This phenomenon would result in high private allelic richness. Several private alleles would then be introduced to the recolonized areas. Therefore, even if only the recolonized area is sampled and compared to areas recolonized from a different refugium, high private allelic richness would be observed. In our results, individuals from the most Western part of the range belong to the light blue cluster, which is well differentiated from the two largest clusters (South Alps grey and Central Eastern Europe orange). However, the low private allelic richness of this South Western and Central France cluster does not seem compatible with the hypothesis of an Iberian refugium origin or with the hypothesis of a French microrefugium. Furthermore, the low allelic richness is also inconsistent with the hypothesis of a secondary contact zone, where a high allelic richness is expected.

Another possibility is to consider a more recent colonization of South Western and Central France by founders originating from Eastern France, South Alps, or Central Eastern Europe and not from unsampled regions. As explained above, the low private allelic richness of the genetic cluster found in South Western and Central France (light blue) seems incompatible with different origins of these individuals. The hypothesis of Klášterský (1968) proposes human-mediated dispersal as the factor responsible for the colonization of South Western and Central France while a natural dispersal may also be responsible for this colonization. Depending on the number of founders, human-mediated secondary introduction as well as natural dispersion could represent a genetic bottleneck, inducing genetic drift. Genetic drift increases population differentiation (Wright, 1931). It also affects allele frequencies, leading to allele fixation or loss, and results in decreased heterozygosity and, to a greater extent, decreased allelic richness (Greenbaum et al., 2014). Therefore, under the hypothesis of Klášterský (1968), we would expect individuals in the Western part of the range to be differentiated from those in the South Alps and Central Eastern Europe due to founder effect. These Western individuals would also exhibit relatively low heterozygosity and allelic richness. Our results are consistent with this expectation regarding genetic differentiation and allelic richness, as the light blue cluster is well differentiated and shows low allelic richness although its heterozygosity is not decreased, which may suggest that the demographic event was relatively recent. However, the natural dispersal of a few founders would also result in such founder effect. Therefore, the Klášterský (1968) hypothesis cannot be prioritize against natural dispersal based on our results. Indeed, at first glance, the clonal lineages containing wild-collected individuals at distant sites and in the Western part of the study range may be used in favor of the Klášterský (1968) hypothesis as only humans can transport clonally reproduced *R. gallica* over large distances. Nevertheless, this argument applies only to the sites where such clonal lineages where detected. This human-dispersal could have taken place either before (supporting the Klášterský (1968) hypothesis) or after the establishment of *R. gallica* in the westernmost parts of the range. Under the latter hypothesis, these parts of the range could have been first naturally recolonized by few founders, followed by an even more recent human-mediated dispersal.

Some wild-collected French samples belong to clusters that are only locally spread (i.e., the yellow, green, dark blue, and pink clusters). Their restricted distribution complicates interpretations, but certain hypotheses regarding their origins can be proposed. The yellow cluster predominantly contains cultivated individuals. Thus, the three wild sites with a high ancestry proportion to the yellow cluster might comprise formerly cultivated individuals that have persisted in the wild since their abandonment. The green cluster includes only three individuals that exhibit a proportion of ancestry greater than 80%. This cluster displays low allelic richness but a high private allelic richness. It is highly differentiated from all other clusters. On the PCoA plot that describes the clone-corrected *Rosa* spp. dataset, individuals from this cluster are not located in the core of the *R. gallica* group but rather between the European *Synstylae* group and the *R. gallica* group (Appendix B.A2). Moreover, among the 17 individuals with a 50-80% ancestry proportion to this cluster, five have a suspected interspecific hybrid origin. Thus, individuals from this cluster might exhibit introgression from other *Rosa* species, such as European *Synstylae*. Given that these species are closely related to *R. gallica*, the hybrids are difficult to identify. This could be exacerbated by noise from the ploidy detection methods we employed (omission of 3x and 5x, lower performance at high ploidy levels), allelic errors, and the allele dosage step. The final two groups, dark blue and pink, are quite restricted and primarily consist of individuals from neighboring sites. Hence, they might be closely related. Given that STRUCTURE inference is sensitive to relatedness, the identification of these groups at K = 8 could be an artifact. However, when examining the clustering results at K = 2 (Appendix G) and the split network partitions at K = 8, individuals in the pink group seem to have emerged from hybridization between Western and more eastern *R. gallica*. Intriguingly, at K = 2, individuals of the dark blue cluster are grouped with those found in the South Alps and Central Eastern Europe, while other individuals from South Western France (i.e. individuals belonging to the light blue cluster) are grouped with cultivated individuals. Therefore, South Western France might have been colonized through several introduction events (either natural or human-mediated).

### Evidences of human-mediated dispersal in France

Boulenger (1932) stated that cultivated *R. gallica* specimens sometimes escaped from cultivation in France. For a particular site, the indication of ancient cultivation could be the inclusion of an individual from the site in the same clonal lineage as a known botanical or cultivated individual. However, the clonal relationship could also be explained by the fact that an individual from the site, harboring interesting traits, has been extracted from wild to be cultivated. Therefore, when individuals from only one site are found in the same clonal lineage as one or several botanical or cultivated individuals, we can not confidently state that the site was a former cultivation place. When multiple sites belong to the same clonal lineage as one or several botanical or cultivated individuals, we can conclude that among the multiple sites, there are ancient cultivation areas. Nevertheless, these sites may or may not include a hypothetical site of origin of a variety directly extracted from the wild. Here, we found multiple sites with individuals belonging to the same clonal lineage as botanical or cultivated individuals. Some of the clonal lineages grouped individuals sampled in more than one site, supporting the hypothesis of Boulenger (1932) which suggests that some *R. gallica* wild sites are certainly ancient cultivations. However, based on our analyses, we can not determine for each site if it is an ancient cultivation or if it is the wild site of origin of a variety. The botanical or cultivated individuals detected in this type of clonal lineages are often of unknown origins, reinforcing the doubt on their origins. Under the hypothesis of Boulenger (1932), the detection of clonal lineages containing wild and non-wild samples is highly dependent on the presence of the clonal non wild specimens in the dataset. If the actual botanical or cultivated variety is not included in the dataset, finding clonal lineages grouping individuals from different sites would be an indication of ancient cultivation. This case was found in Spain, making the ancient cultivation hypothesis possible for the clonal lineage J. This would also be in accordance with the hypothesis of a secondary origin of Spanish populations as written by Klášterský (1968). A similar grouping was found for three individuals collected from three different sites in Slovenia. These were sampled in the botanical garden of Ljubljana from three plants that were reported to be collected in three distant wild Slovenian sites. Therefore, an explanation for the clonal relationship would be plant identification loss or mistake during transplantation. Additionally, we found clonal lineages gathering botanical or cultivated individuals and individuals from two sites in Bosnia and Herzegovina. These individuals from Bosnia and Herzegovina were considered as wild for the analyses but were collected in gardens, therefore, the found clonality supports their non wild nature.

### Conclusion and perspectives

This study provides the first near range-wide characterization of the genetic diversity of *R. gallica*. Our results reveal that both natural expansion and human-mediated dispersal have shaped the evolutionary history of the species. Natural postglacial expansion appears to have been the main driver of colonization for populations in the South Alps and Central Eastern Europe, whereas the most western populations were likely established more recently through either natural dispersal with founder effects, human-mediated dispersal, or a combination of both.

Focusing on France, we uncovered a complex genetic structure suggesting contrasting demographic histories between the eastern and western parts of the country. Eastern France may have been recolonized following the same routes as the South Alps and Central Europe, while central and western France appear to have been recolonized more recently, potentially through multiple dispersal events.

While our results allow us to formulate hypotheses regarding the origin of these populations, several questions remain unresolved, in particular the origin of the western populations and the potential contribution of southern refugia. Progressing from descriptive to hypothesis-testing approaches would provide stronger evidence to disentangle these scenarios. The Approximate Bayesian Computation (ABC) framework (Beaumont et al., 2002) offers a powerful and flexible tool to compare alternative demographic models. In a coalescent framework, haplotypes are simulated rather than diploid or polyploid genotypes, meaning that one needs to simulate the number of haploid genomes corresponding to the sampled allele copies. The main challenge lies in how these haplotypes are combined into genotypes for genotype-level summary statics: assumptions about inheritance mode will influence expected genotype frequencies and heterozygosity. Future work should therefore evaluate the impact of alternative inheritance models on summary statistics before applying ABC-based inference to polyploid populations.

Finally, the difficulty in interpreting clonal relationships between wild and cultivated individuals (distinction between cases of ferality or remarkable wild individuals put into cultivation) shows the need for a better understanding of *R. gallica* domestication: do cultivated *R. gallica* have a single origin from wild *R. gallica* or have they benefited from successive introgressions from the wild pool?

## Appendices

**Appendix A.**
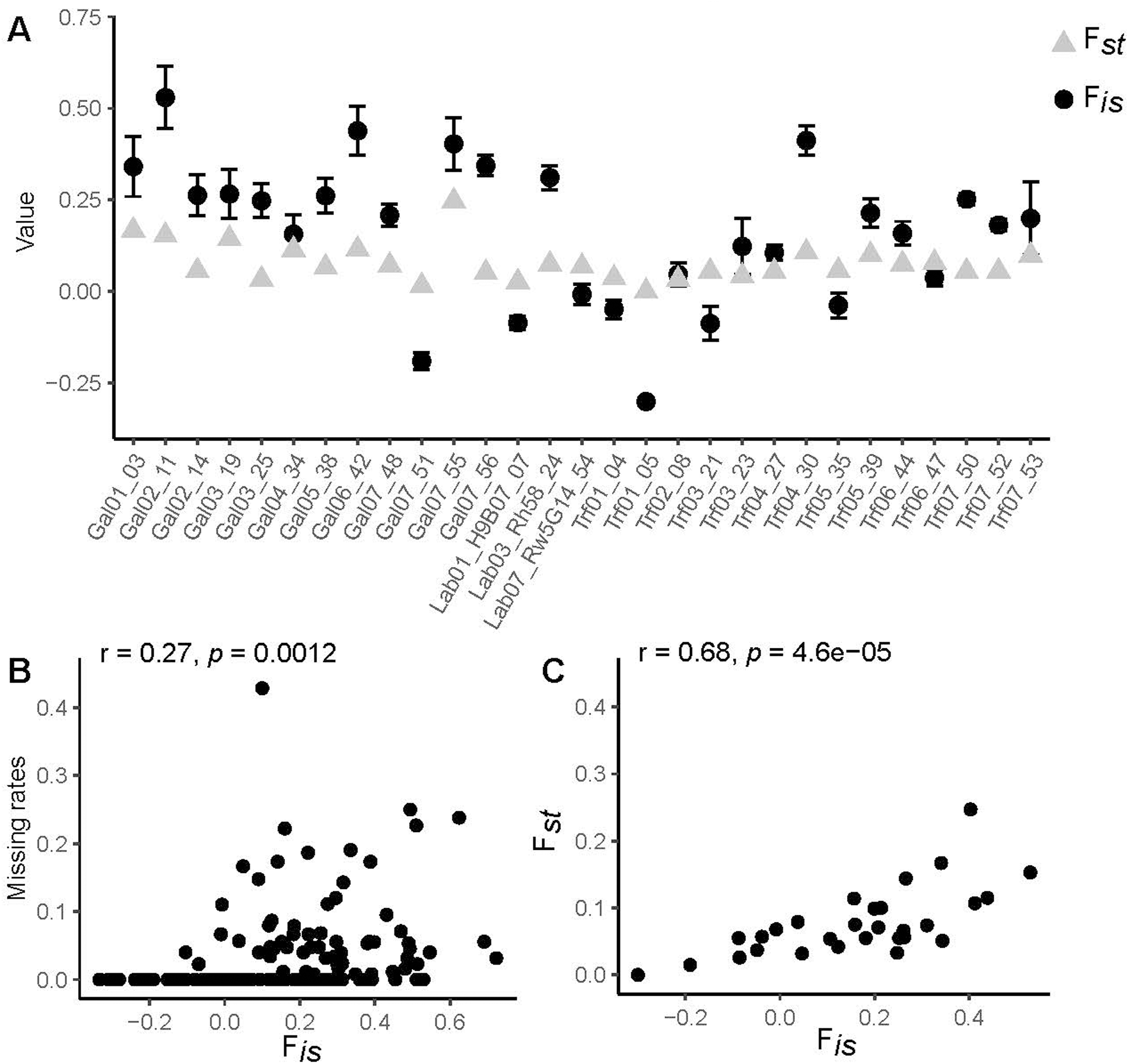
Exploration of null alleles. (A) F*_is_* and F*_st_* values per locus for the 29 markers, calculated for the five STRUCTURE clusters with more than 20 non-admixed individuals. F*_is_* standard errors were estimated using jackknifing over clusters. (B) Pearson’s correlation between missing rates and F*_is_* in each cluster. (C) Pearson’s correlation between F*_st_* and F*_is_* averaged over clusters.

**Appendix B.**
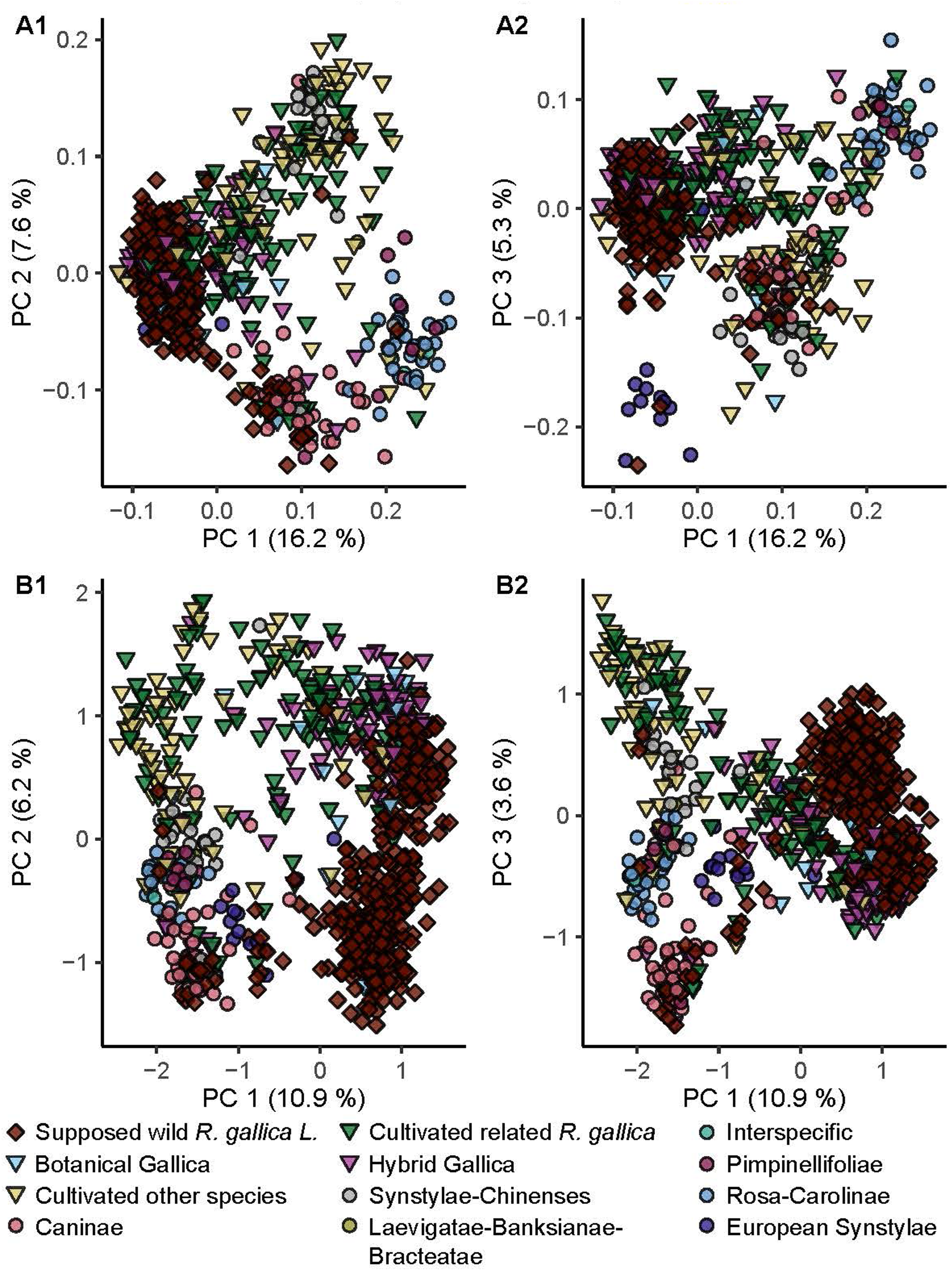
Multivariate analyses describing the *Rosa* spp. dataset. (A) Principal Coordinates Analysis based on pairwise k-mer dissimilarities. (B) Principal Component Analysis of intra-individual allele frequencies. A1 and B1 correspond to individual values for the first and the second principal components while A2 and B2 show the first and the third principal components whatever the multivariate analysis used. Cultivated individuals are represented as triangles, whereas wild *R. gallica* are shown as diamonds, and other *Rosa spp.* individuals are represented by circles. Colors correspond to the horticultural or botanical groups. Botanical sections are regrouped according to Debray et al. (2022).

Comments on Appendix B: To assess the reliability of our k-mer based method for dissimilarities calculation, we compared the PCoA based on these dissimilarities and the PCA based on intra-individual allele frequencies performed on the clone corrected *Rosa* spp. dataset. Both analyses separated most of the *R. gallica* specimens from specimens belonging to other species. In details, on the PCoA plots, most of the *R. gallica* specimens were grouped at the left of the plot when visualizing both the axis one and two (A1, dark red diamonds), and the axis one and three (A2, dark red diamonds). For the PCA, most of the *R. gallica* were located at the right of the plot when visualizing the axis one and two and the axis one and three (B1-B2, dark red diamonds). Comparing the k-mer dissimilarity based PCoA and the PCA, we saw that specimens tended to be clustered according to their botanical section. Indeed, on the PCoA, the European *Synstylae* were grouped apart at the bottom left of the subfigure A2 (purple circles). They also grouped together on the PCA (B1-B2, purple circles). The *Caninae* specimens formed a group at the bottom of the subfigures A1, B1 and B2 (pink circles). The individuals belonging the *Rosa*-*Carolinae* sections clustered together on both the PCoA and PCA. On the PCoA, they were located on the right side of the two plots (A1-A2, blue circles) and on the PCA, these individuals were located at the middle left of the two plots (B1-B2, blue circles). Individuals belonging to the sections *Synstylae*-*Chinenses* were grouped with some cultivated specimens, related or not to *R. gallica* (red and yellow triangles), at the top of the subfigure A1 (PCoA, grey circles). The *Synstylae* or *Chinenses* were also grouped together on the PCA (B1-B2, middle left, grey circles). Looking at the separation between the groups identified above, the *Synstylae*-*Chinenses* group was more distinguishable from the group of *Rosa*-*Carolinae* on the k-mer based PCoA (A1, right top and middle right) than on the allele frequencies based PCA (B1-B2, both groups were located close to each other in middle left). Several suspected *R. gallica* specimens were not included into the major group of individuals of this species. Indeed, a few *R. gallica* (dark red diamonds) were grouped with *Caninae* (pink circles) whatever the method used. Additionally, few other suspected *R. gallica* individuals were found in the groups of *Rosa*-*Carolinae*, *Synstylae*-*Chinenses* and European *Synstylae*. The location of these individuals suggested a mis-identification. Finally, the grouping obtained using either a PCoA based on k-mer dissimilarities or PCA based on intra-individual allele frequencies were globally similar. Moreover, the percentages of explanation were of the same order for both methods.

**Appendix C.**
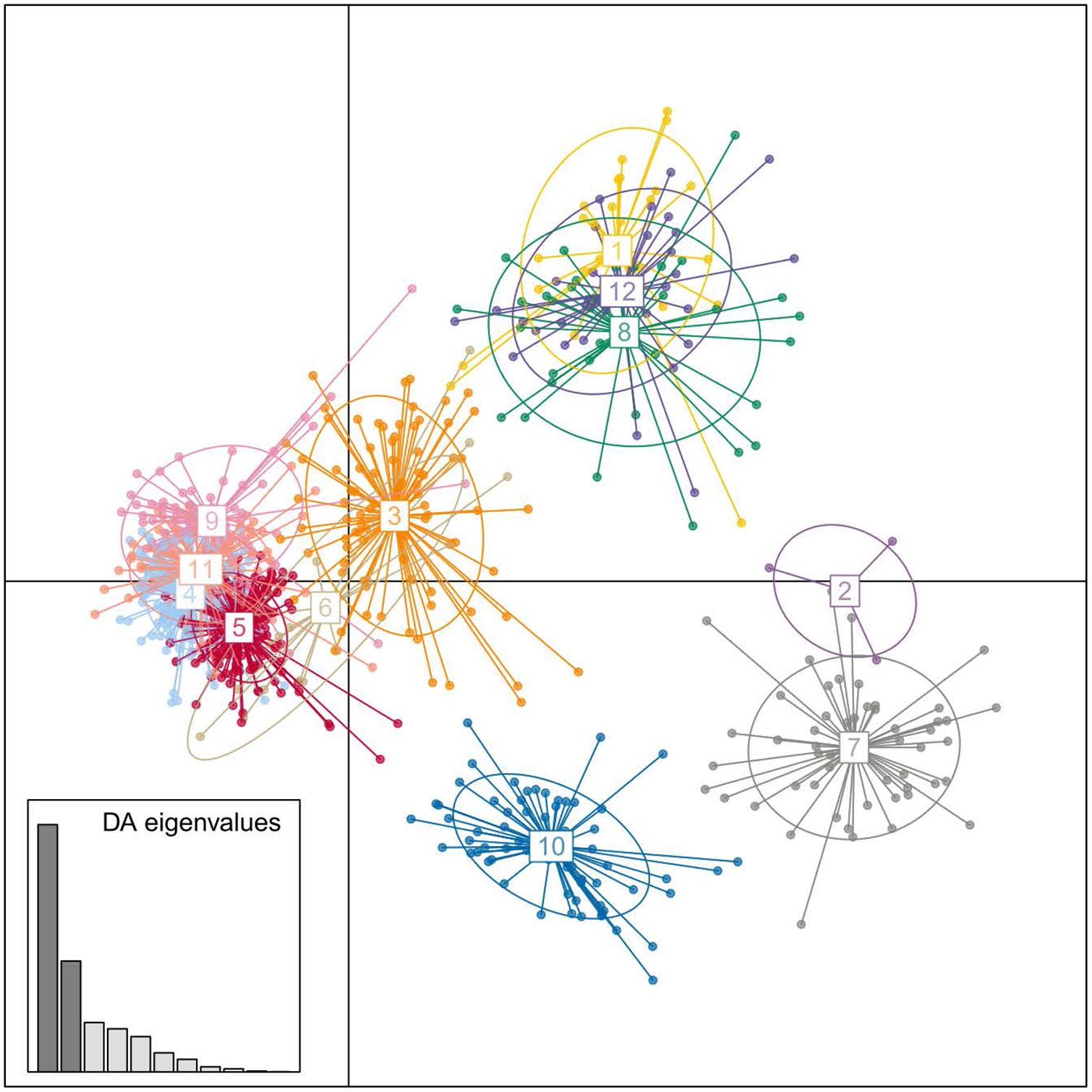
Scatterplot of the first two DAPC discriminant functions. The DAPC was done on the clone-corrected *Rosa* spp. dataset setting the number of genetic groups to 12. Colors indicate genetic group memberships.

**Appendix D.**
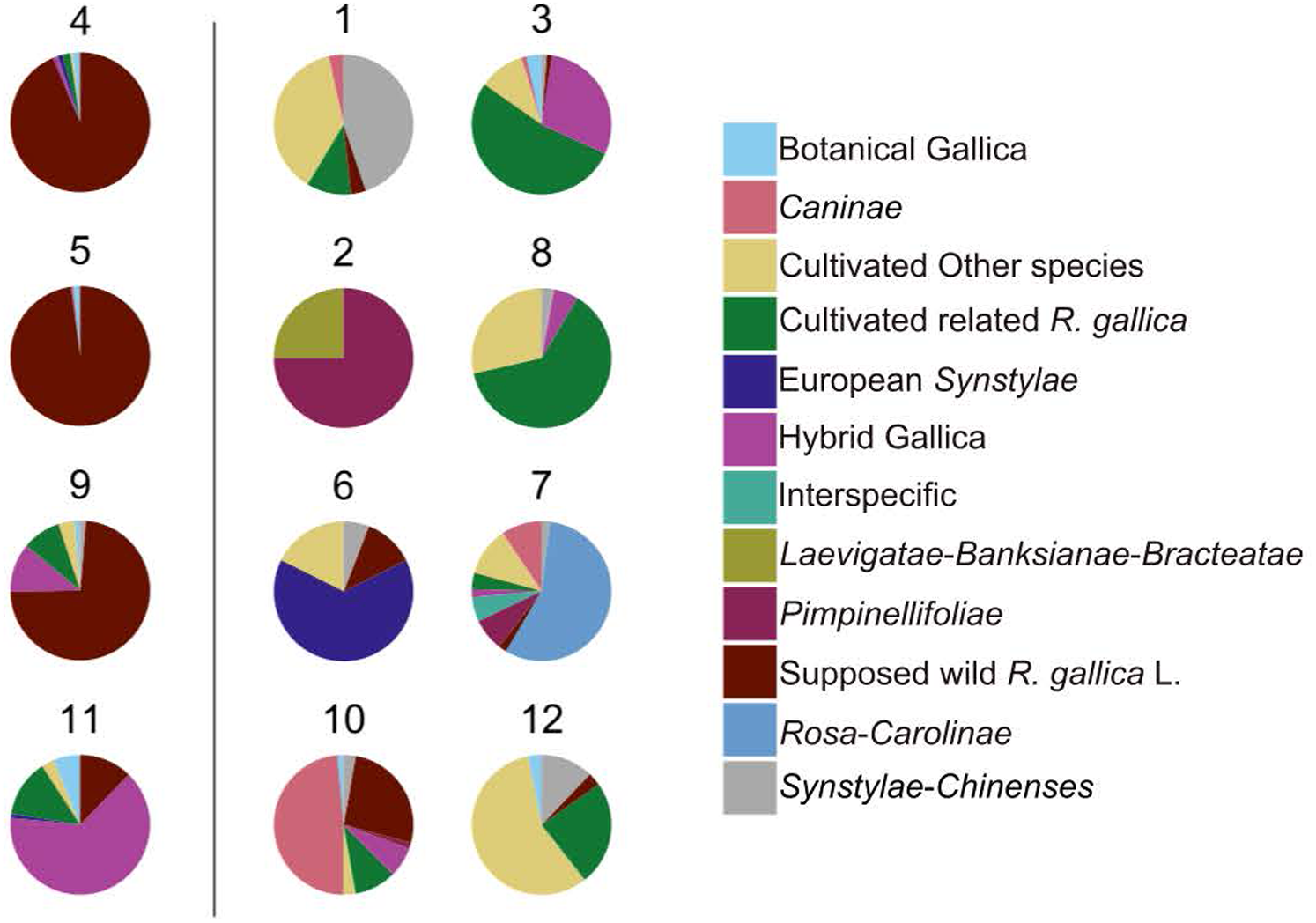
Content of the genetic groups obtained by performing a DAPC on the clone-corrected *Rosa* spp. dataset. Group content is detailed in term of botanical sections (for wild specimens) and horticultural groups (for cultivated specimens). Some horticultural groups and botanical sections were merged to improve figure clarity. In detail, Damasks, Albas, Mosses, Portland, Centifolia and Bourbons were regrouped under the name “Cultivated related *R. gallica*”, cultivars belonging to other horticultural groups (except Hybrid Gallicas) were grouped together under the name “Cultivated other species”. Group assignation used for this figure are available in Table S1. The genetic groups 4, 5, 9 and 11 were considered as groups of *R. gallica* individuals.

**Appendix E.**
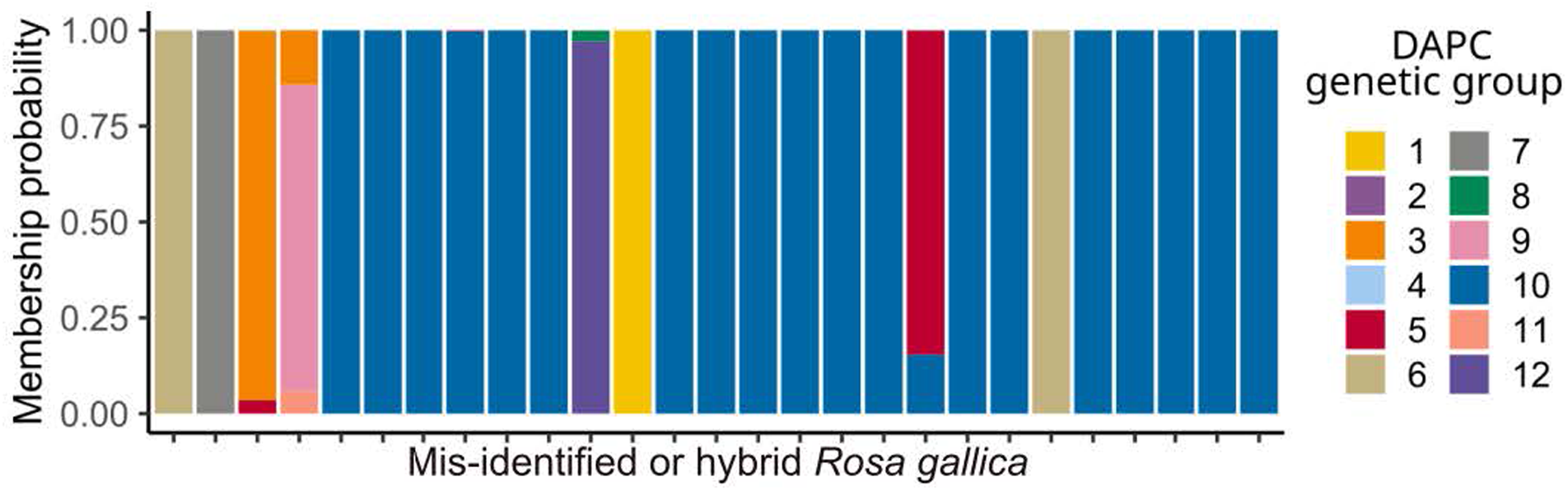
Compoplot of the DAPC membership probabilities of the 27 misidentified *R. gallica* or hybrids with other *Rosa* species. DAPC genetic groups content are available in Appendix D. For instance, 19 individuals show high membership probability to the DAPC genetic group 10 that contains a majority of *Caninae* specimens.

**Appendix F.**
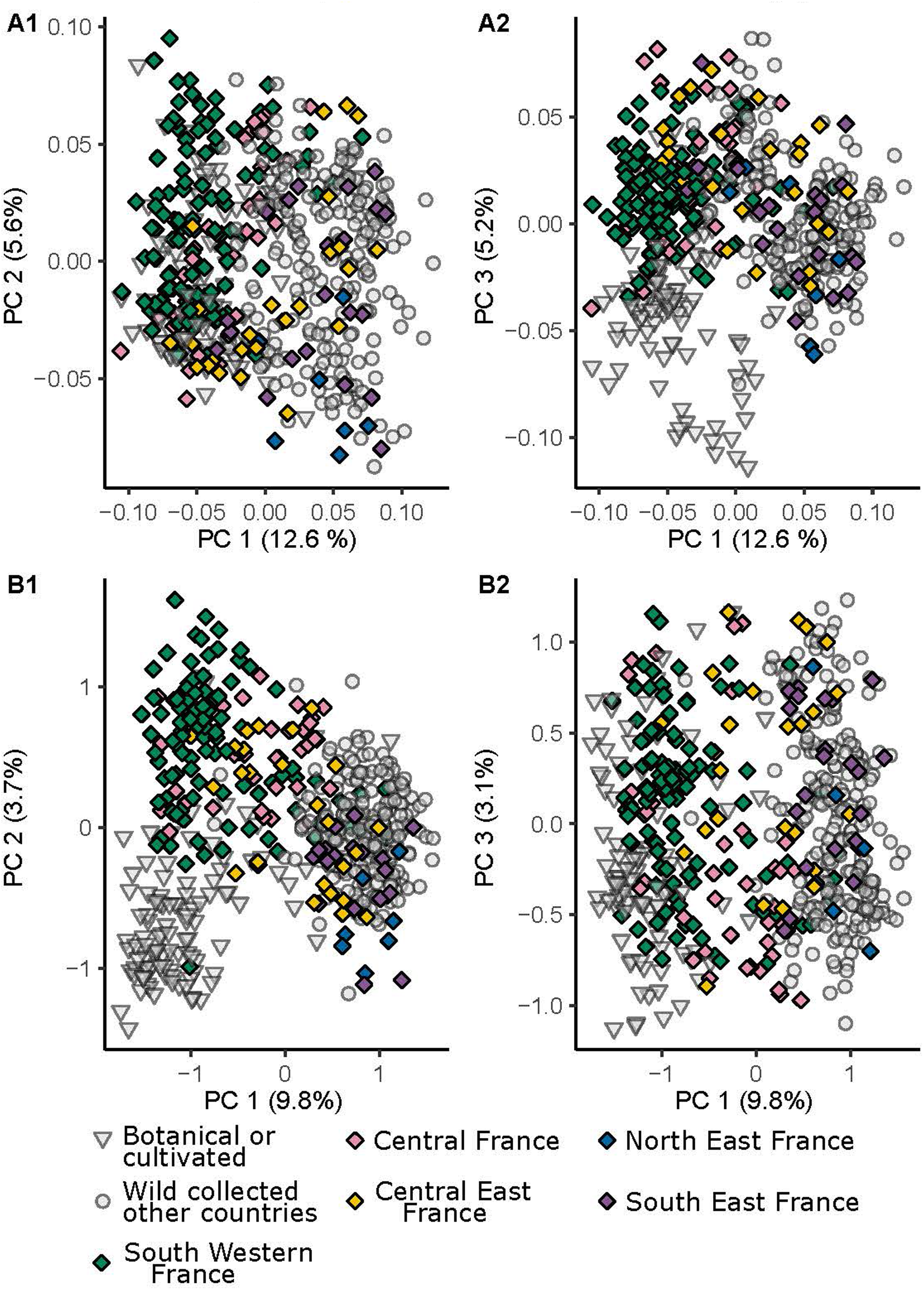
Multivariate analyses describing the *R. gallica* dataset. (A) PCoA based on pairwise k-mer dissimilarities. (B) PCA based on intra-individual allele frequencies. A1 and B1 correspond to individual coordinates for the first and the second principal components while A2 and B2 show the first and the third principal components whatever the multivariate analysis. Cultivated or botanical individuals are depicted as triangles, whereas French wild-collected ones are shown as diamonds, and other wild-collected individuals are represented by circles. Colors correspond to the French regional groups inferred from community detection analysis (Fig. 3). Non French or non wild individuals are in grey.

**Appendix G.**
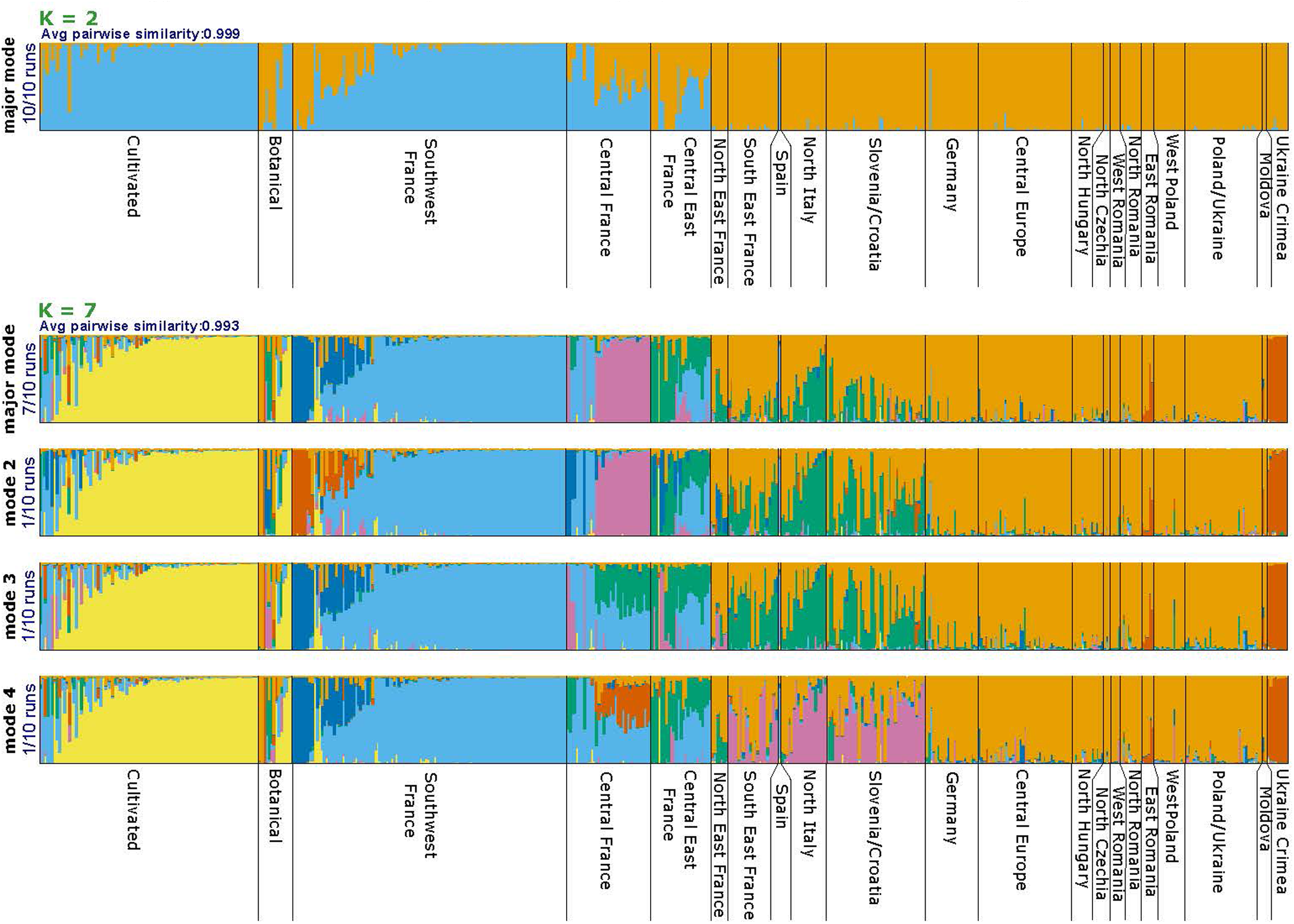
STRUCTURE bar plots of the ancestry proportions at K = 2 and K = 7 for the *R. gallica dataset*

## Acknowledgements

We thank the following contributors for their assistance with the collection and wet-lab analyses of wild and cultivated rose samples: P. Bardin, O. Bezsmertha, F. Bonali, J. Chameau, E. Chancerel, Z. Compagnie, F. G. Dunkel, O. Elsner, A. Fedorova, M. Fickel, C. Fy, C. Gilli, D. Gomez, E. Guichoux, E. Hauser, P. Heitzler, J. Herbst, M. Hohla, G. Kleesadl, L. Lambelin, A. Lugmair, C. Marie, V. Martínez Francés, J. Marie-Magdelaine, L. Meierott, G. Michel, J. Mondion, G. F. Haefner, W. Mucher, D. Pastor, U. Raabe, S. Rios, A. Rosenbauer, W. K. and M. Rottensteiner, J. Schach, C. N. Schröder, H. Tinguy, E. Trummer-Fink and V. Wisseman. We also thank the Muséum national d’Histoire naturelle (MNHN) for access to its collection, as well as the citizen science platform “Les Herbonautes” (MNHN/RECOLNAT; ANR-11-INBS-0004) for providing data associated with herbarium samples. We further thank the Conservatoires botaniques nationaux, in particular the CBNPMP, CBNBP, CBNSA, CBNA, CBNMC, CBNMed and the CBN Brest for their assistance in locating and collecting wild *R. gallica* individuals. We are grateful to the Loubert Rose Garden, La Cour de Commer rose garden (Cour-de-Commer, France), Jumaju rose garden (Montchamp, France), Jardin botanique de la Tête d’Or (Lyon, France), Désert rose garden (Bouzon Gellenave, France), Grande roseraie de Lyon, La Beaujoire rose garden (Nantes, France), Arboretum des Barres (Nogent-sur-Vernisson, France), the botanical garden of the University of Angers pharmaceutical faculty (France) and the Norwegian Arboretum rosarium of the university of Bergen (Norway) for providing access to their collections of cultivated roses. We also acknowledge the Pome Fruits and Rose BRC for providing DNA and associated data for the studied accessions. The authors thank UE HORTI, INRAE, Horticulture Experimental Facility (doi:10.15454/1.5573931618268674E12) for cultivation of some of the studied rose accessions. Microsatellite development and sequencing were carried out at the PGTB (doi:10.15454/1.5572396583599417E12) with the assistance of B. Tyssandier and A. Delcamp. The authors are also grateful for the technical support provided by the ANAN platform of the SFR QUASAV including M. Bahut, during DNA extractions. Operations of sampling leaves of French *R. gallica* individuals were carried out under prefectural licenses, as the species is protected. The authors are grateful for these authorizations. Finally, we thank F. Jabbour, R. Bacilieri, and V. Wissemann for their advices given throughout the PhD thesis of C. Pawula.

## Data, script, code, and supplementary information availability

Sequencing data of the samples used in the analyses are available under ENA BioProject PRJEB105314. Sample metadata are provided as supplementary information and are also available in an analysis-ready format at https://doi.org/10.5281/zenodo.14686589 (Pawula et al., 2026). Scripts and codes are available online: https://doi.org/10.5281/zenodo.14686589 (Pawula et al., 2026). Supplementary information is available online: https://doi.org/10.5281/zenodo.14686589 (Pawula et al., 2026).

