## Supplementary information for "Range wide analysis of genetic diversity and structure gives insights into *Rosa gallica* L. evolutionary history"

This document presents the supplementary Figures and Tables associated with "Range wide analysis of genetic diversity and structure gives insights into *Rosa gallica* L. evolutionary history". All supplementary figures are included here, whereas some large tables are provided as separate files available at `https://doi.org/10.5281/zenodo.14686589`.

This document also describes the Codes and Scripts used in the present study, available at `https://doi.org/10.5281/zenodo.14686589` in the folder **rosa_gallica_genpop**. Scripts also includes instruction for the reproduction of analyses done with standalone software from the Graphical User Interface. To ease the reproduction by skipping time consuming step, important intermediate data files are already available in sub-folders. Please start by reading the README.html file available in the **rosa_gallica_genpop** if you need details on each file content. A copy of the README content is available in section 4

### 1 Overview

#### Supplementary Figures

- **Figure S1: Pairwise euclidean distance distribution used for the clone detection.**
- **Figure S2: Comparison of different $K$ values for k-means clustering using Bayesian Information Criterion.**
- **Figure S3: Distribution of DAPC membership probability at $K = 12$ for the *Rosa* sp. dataset.**
- **Figure S4: Distribution of the maximal ancestry percentages inferred using STRUCTURE at $K = 8$ on the *R. gallica* dataset.**
- **Figure S5: Physical map of the chromosome locations of the 29 loci.**
- **Figure S6: Determination of optimal K values for STRUCTURE inference.**
- **Figure S7: Exact mapping of STRUCTURE mean ancestry proportions per site of collection at $K = 8$.**

#### Supplementary Tables

- **Table S1: Individual metadata.** Available as Standalone. Columns are described in the present document.
- **Table S2: Sample metadata.** Available as Standalone. Columns are described in the present document.
- **Table S3: Primer pairs sequence and location.** Available as Standalone. Columns are described in the present document.
- **Table S4: Final parameters of the allele calling pipeline.** Available as Standalone. Columns are described in the present document.

- Table S5: **Ancestry proportions of *R. gallica* dataset samples inferred by STRUCTURE at $K = 8$ for the major mode.** Available as Standalone. Columns are described in the present document.
- Table S6: **Number of clonal lineages and sites per geographical regions.** Available in the present document.
- Table S7: **Ploidy estimation results and FDSTools ploidy parameters.** Available as Standalone. Columns are described in the present document.
- Table S8: **Quality and polymorphism of the 29 loci, assessed on the *Rosa gallica* dataset.** Available in the present document.
- Table S9: **DAPC cluster membership probabilities of the *R. gallica* dataset samples at $K = 12$ for the major mode.** Available as Standalone. Columns are described in the present document.
- Table S10: **Pairwise $\rho$ differentiation between geographical regions**. Available in the present document.
- Table S11: **Private allelic richness per geographical region**. Available in the present document.

#### 2 Supplementary Figures

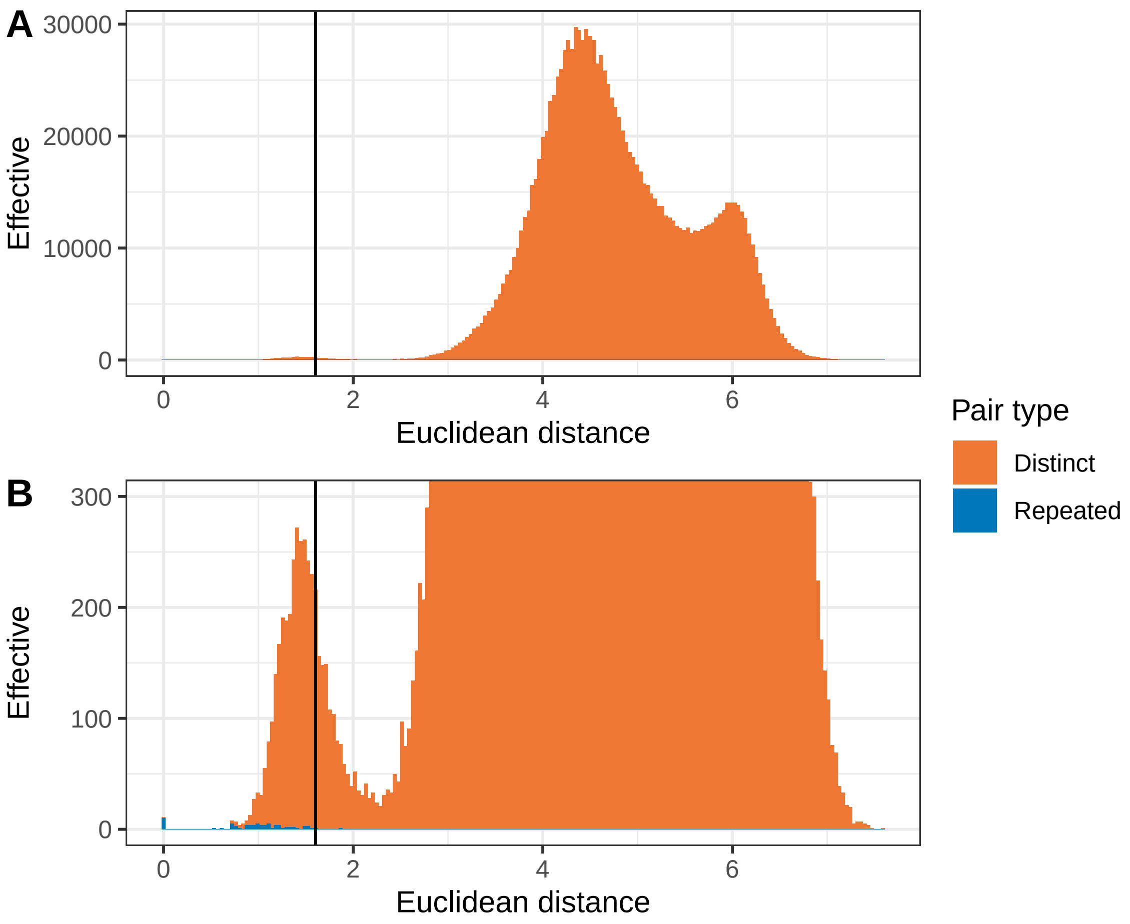


Figure S1: Pairwise euclidean distance distribution used for the clone detection. (A) correspond to the full distribution view while (B) is a focused view. The vertical black line represent the threshold of 1.603 used. Colors indicate whether the pairwise distance was calculated between known repeated or distinct individuals.

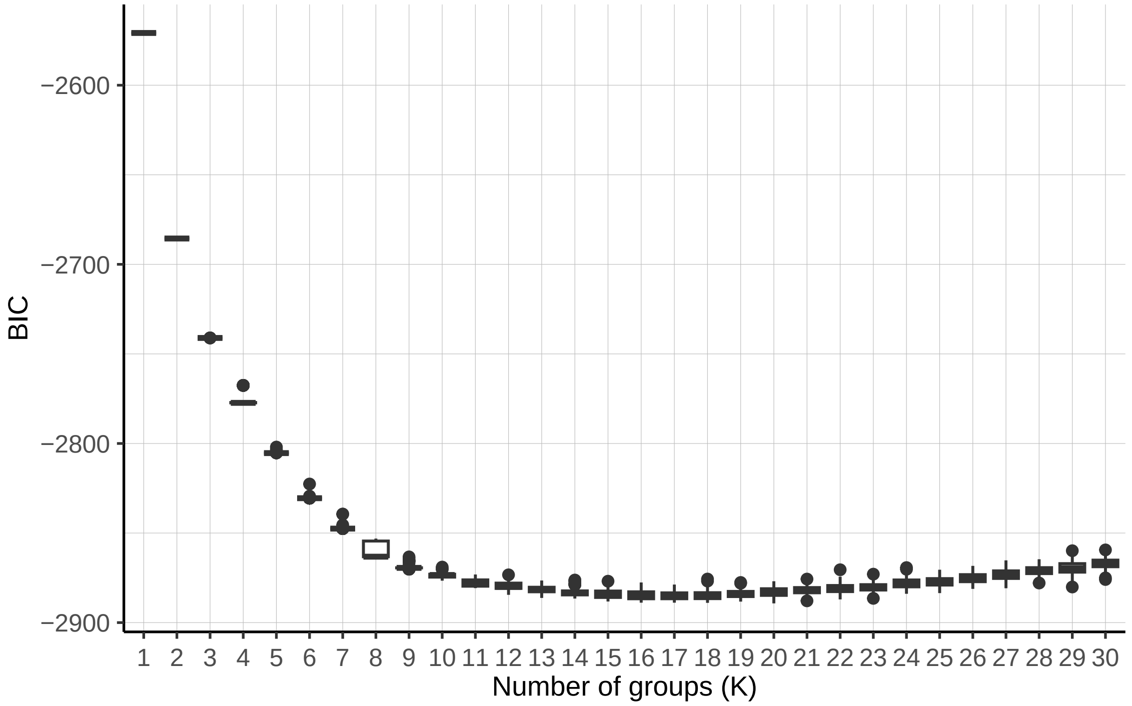


Figure S2: Boxplots of Bayesian Information Criterion (BIC) for different values of $K$ in k-means clustering analyses of the *Rosa* sp. dataset. To get these results, the function find.clusters from the R packages adegenet v2.1.10 was run independently 100 times on the *Rosa* sp. dataset. An optimal value of $K = 12$ was selected as the BIC did not significantly decrease for higher $K$ values.

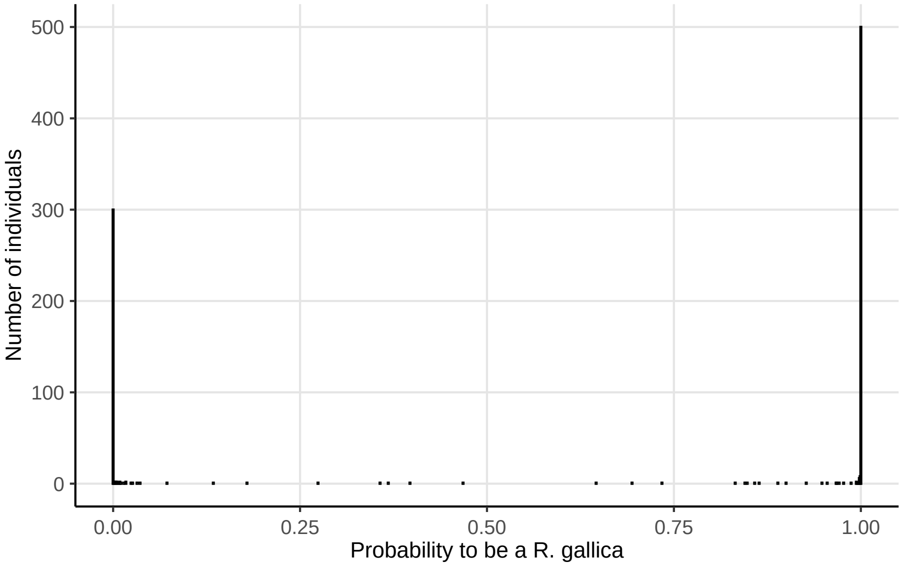


Figure S3: Distribution of DAPC membership probability at $K = 12$ for the *Rosa* sp. dataset. Due to this nearly 0/1 distribution, the threshold chosen to discriminate between admixed and non-admixed individuals was set to 0.99.

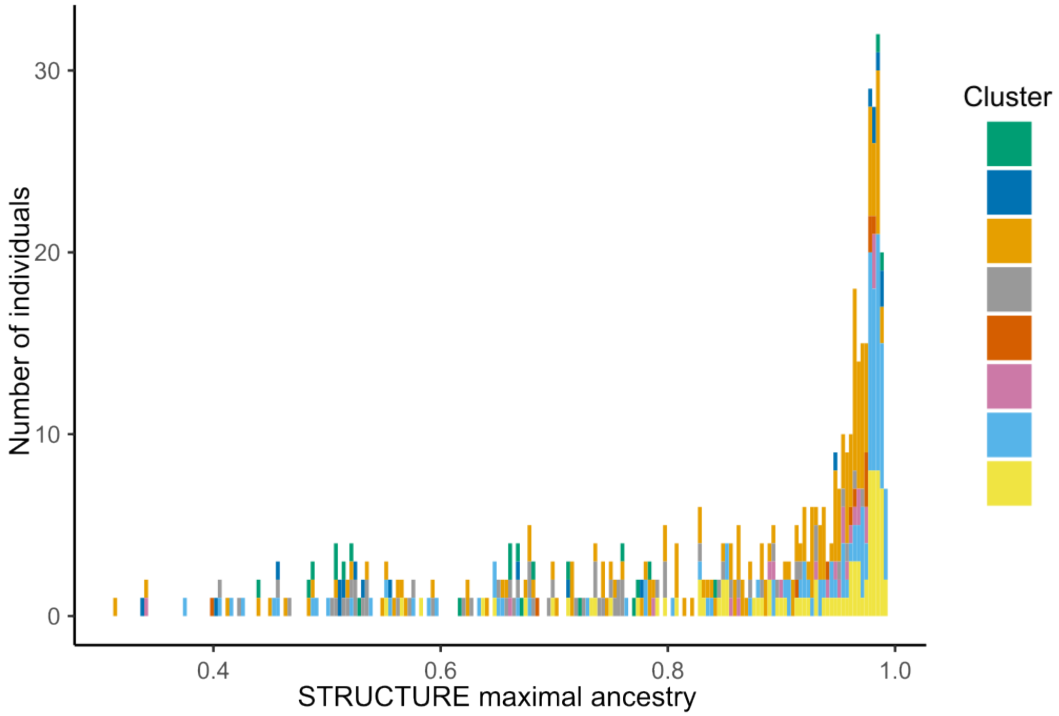


Figure S4: Stacked barplots of the maximal ancestry percentage per individual inferred using STRUCTURE at $K = 8$ on the *R. gallica* dataset. The threshold chosen to discriminate admixed and non admixed was set to 0.80. Associated data, can be found in Table S5

.

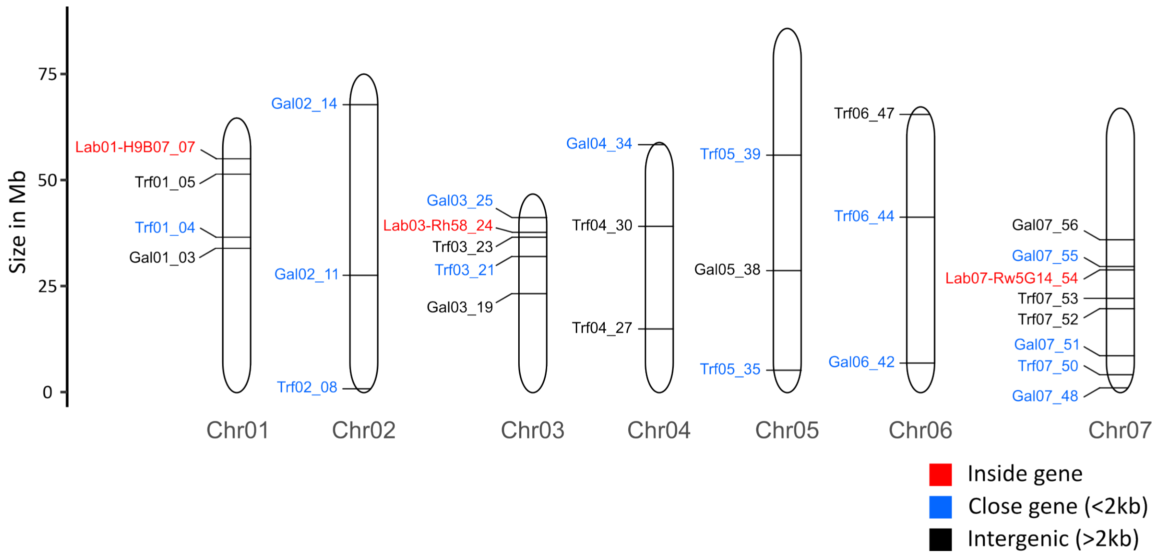


Figure S5: Physical map of the chromosome locations of the 29 loci. Gene proximity is indicated by colours. Loci names are colored in red if the locus is located within a gene, in blue if a gene is nearby (at most 2 kb), and in black if the closest gene is at least 2 kb away from the locus.

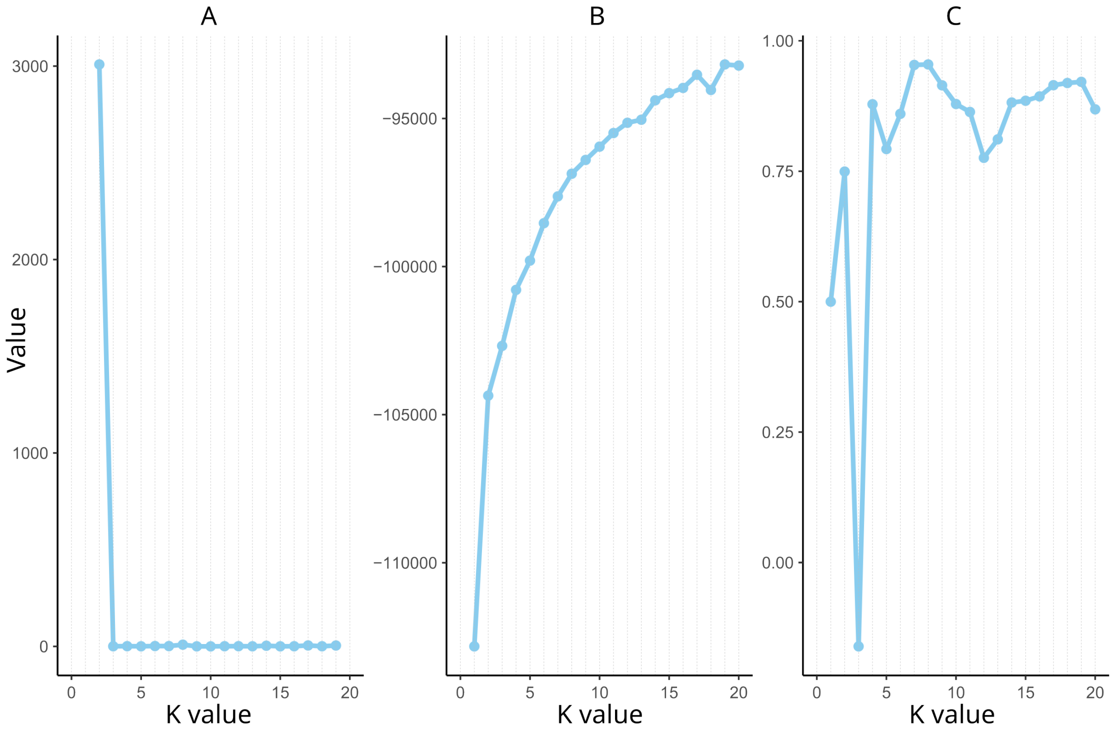


Figure S6: Determination of optimal K values for STRUCTURE inference of the *R. gallica* dataset Three estimators were computed: (A) the $\Delta K$ (Evanno et al., 2005), (B) the $P(X|K)$ (Pritchard et al., 2000) and (C) the Parsimony Index (Wang, 2019). Inference was done on the *R. gallica* dataset.

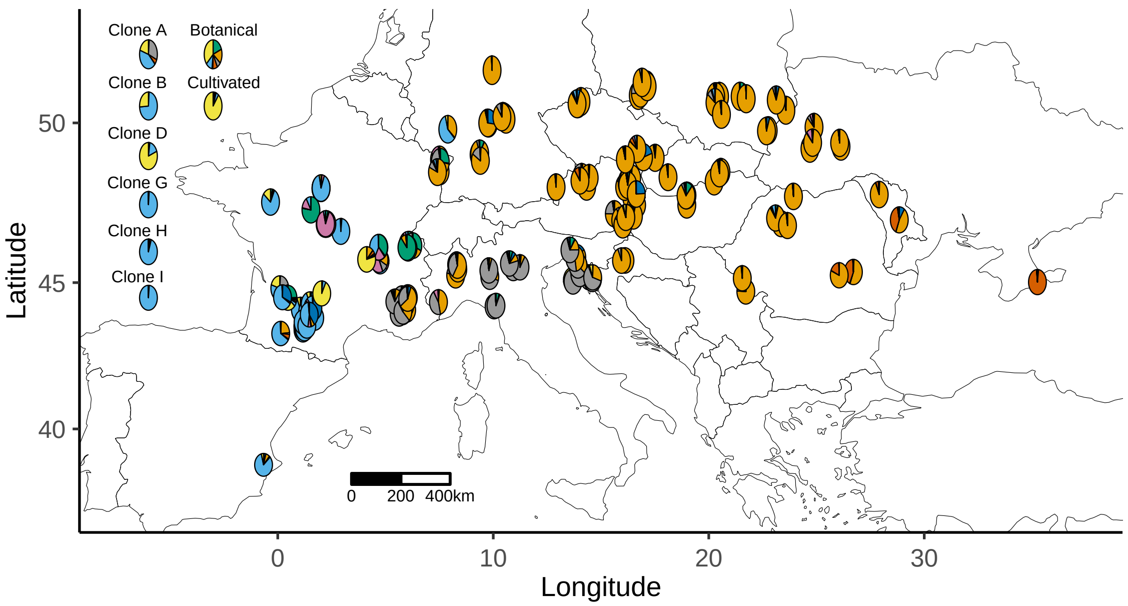


Figure S7: Exact mapping of STRUCTURE mean ancestry proportions per site of collection at $K = 8$. Ancestry proportions of the representative individual of the clonal lineages A, B, D, G, H and I, as well as mean ancestry proportions of cultivated and botanical individuals are also shown on the left of the map.

### 3 Supplementary Tables

**Table S1:** Individual metadata. Available as Standalone.

The table is available as a comma separated value CSV file at https://doi.org/10.5281/zenodo.14686589. This dataframe contains 1586 rows and 14 columns
The columns are:

- **Individual_ID**: Name of the individual.
- **Species**: Species of the specimens. When available, the botanical variety is specified. If species identification was not possible, the botanical section is indicated instead.
- **Class**: Individual classification as botanical or cultivated, Specimens from other *Rosa* sp., wild *R. gallica*. The full-sib and half-sib used for clone detection test are also indicated.
- **Hort_group_Bot_section**: The horticultural group of cultivated individuals and the botanical section of wild individuals. Horticultural groups are named according Cairns (2003). Cultivated individuals were affected to horticultural groups according to Liorzou et al. (2016) and/or to the HelpMeFind database (https://www.helpmefind.com/). The *Synstylae* are divided into *Synstylae* and European *Synstylae* according Debray et al. (2022).
- **Group_appendix_D**: Group used in Appendix D.
- **Country_Rgallica**: Country of origin of the wild *R. gallica*.
- **Site**: Name of the site were the *R. gallica* were collected.
- **Lat_trunc**: Truncated latitude in WGS84 of wild *R. gallica*.
- **Long_trunc**: Truncated longitude in WGS84 of wild *R. gallica*.
- **Location_accuracy**: Accuracy of the GPS location, either Site, City or Country.
- **Origin**: Origin of the sample, classified as Wild, Garden, or Scientific material. A few individuals sampled in gardens were considered Wild based on the provider's information.
- **Provider_institution**: When applicable, the institution of the sample provider is indicated.
- **Collection_year**: Year of sampling.
- **Type_of_sample**: Nature of the received sample, which can be either freeze-dried, silica-dried, herbarium specimen, or already extracted DNA.

**Table S2:** Sample metadata. Available as Standalone.

The table is available as a comma separated value CSV file at https://doi.org/10.5281/zenodo.14686589. This dataframe contains 1710 rows and 10 columns
The columns are:

- **Individual_ID**: Name of the individual.
- **Sample_ID**: Code of the sample. As some individuals were genotyped several times, there might be several Sample_ID for the same Individual_ID.
- **FDSTools_dataset**: Samples used for FDSTools parameters optimization, either diploid, tetraploid or synthetic tetraploid samples.
- **Rosa_sp_dataset**: Samples included in the *Rosa* sp. dataset.
- **Clonal_group**: Clone detection results. The number corresponds to the clonal lineage to which the sample belongs.
- **Inter_Clonal_group**: Clonal lineages classification used in figure 1. For wild specimens "No_inter_clone" indicates that in a given clonal lineage, there are not two wild *R. gallica* from different sites. For cultivated specimens, "No_inter_clone" indicates that there is only cultivated specimens in the clonal lineage.
- **ID_ind_after_clcorr**: For each clonal lineage, the column indicates which sample is retained for the next steps. For clonal lineages A to K, a new name is assigned to the selected representative sample.
- **Clone_corrected_Rosa_sp_dataset**: Samples included in the clone corrected *Rosa* sp. dataset.
- **Rosa_gallica_dataset**: Samples included in the *Rosa gallica* dataset.
- **Region_groups**: Regional group of samples belonging to the *Rosa gallica* dataset inferred using distance network. This column is used in Figure 3, Figure 8, Table S9 and Table S10.

**Table S3:** Primer pairs sequence and location. Available as Standalone.

The table is available as a comma separated value CSV file at https://doi.org/10.5281/zenodo.14686589. This data frame contains 60 rows and 9 columns.
The columns are:

- **Locus**: Name of the Locus.
- **chrom**: Chromosome of the locus.
- **chromStart**: Locus starting location.
- **chromEnd**: Locus ending location.
- **WORD_TRANS**: Repeat unit.
- **PRIMER_LEFT_SEQUENCE**: Left primer sequence.
- **PRIMER_RIGHT_SEQUENCE**: Right primer sequence.
- **Reference genome access**: Link to access the reference genome.
- **Source**: Source of the marker design.

**Table S4:** Final parameters of the allele calling pipeline for the 60 loci. Available as Standalone.

The pipeline was designed by Lepais et al. (2020) based on the FDSTools v1.2 analysis toolkit (Hoogenboom et al., 2017). The table is available as a comma separated value CSV file at https://doi.org/10.5281/zenodo.14686589. This data frame contains 60 rows and 10 columns. The columns are:

- **Locus**: Name of the Locus.
- **Calling_strategy**: The focusing strategy, either RepeatFocused (ignoring flanking sequences for the calling) or Full-length (taking into account the whole sequence for the calling).
- **Parameter_set**: Set of parameter, either parameter set 1 or 2. The parameters are then detailed in the columns "s-1","s+1","m","M" and "x".
- **s-1**: A *Stuttermark* parameter (included in FDSTools). It corresponds to the relative maximum read coverage of the n-1 repeat stutter compared to an allele with n repeats to consider it as a stutter, otherwise, it is considered as another allele.
- **s+1**: A *Stuttermark* parameter (included in FDSTools). It corresponds to the relative maximum read coverage of the n+1 repeat stutter compared to an allele with n repeats to consider it as a stutter, otherwise, it is considered as another allele.
- **m**: An *AlleleFinder* parameter (included in FDSTools). It corresponds to the relative maximum read coverage of an allele compared to the read coverage of the allele with the highest read coverage to consider it as a true allele, otherwise, it is considered an error.
- **n**: An *AlleleFinder* parameter (included in FDSTools). Minimal read coverage of the most covered allele, otherwise, the genotype is not determined, and the locus genotype is classified as missing.
- **M**: An *AlleleFinder* parameter (included in FDSTools). If, at a locus, an individual has a supernumerary (beyond the ploidy level) allele with a read coverage (relative to the most covered allele) exceeding this threshold, the locus is classified as missing due to potential contamination or its paralogous nature.
- **x**: An *AlleleFinder* parameter (included in FDSTools). It corresponds to the maximal number of locus allowed to be detected as contaminated or paralogous, otherwise, the whole multilocus genotype is set as missing.
- **A**: An *AlleleFinder* parameter (included in FDSTools). Ploidy class of the individual either 2, 4 or 6.

**Table S5:** Ancestry proportions of *R. gallica* dataset samples inferred by STRUCTURE at $K = 8$ for the major mode. Available as Standalone.

The table is available as a comma separated value CSV file at: https://doi.org/10.5281/zenodo.14686589. This data frame contains 519 rows and 13 columns.
The columns are:

- **Sample_ID**: Name of the sample.
- **ID_ind_after_clcorr**: For some samples, the new name assigned to the selected representative sample of clonal lineages, cf Table S2.
- **Cluster1-Green** to **Cluster8-Yellow**: Ancestry proportions to the corresponding clusters inferred by STRUCTURE at $K = 8$.
- **Cluster_Max**: For each individual, the cluster showing the maximum ancestry.
- **Max_ancestry**: For each individual, the ancestry proportion to the corresponding Cluster_Max.
- **Individual_Admixture**: Individual admixture based on the 0.80 Max_ancestry threshold.

**Table S6:** Number of clonal lineages and sites per geographical region.

| Regional Group | Number of clonal lineages | Number of sites |
|---|---|---|
| Slovenia/Croatia | 41 | 16 |
| Central Europe | 39 | 24 |
| Central France | 35 | 12 |
| Central East France | 25 | 11 |
| North East France | 7 | 4 |
| South East France | 21 | 12 |
| South West France | 114 | 39 |
| Germany | 22 | 12 |
| North Hungary | 13 | 8 |
| North Italy | 19 | 12 |
| West Poland | 13 | 5 |
| Poland/Ukraine | 32 | 15 |
| East Romania | 5 | 2 |
| North Romania | 9 | 4 |
| Ukraine Crimea | 9 | 1 |
| Moldova | 2 | 2 |
| North Czechia | 3 | 3 |
| Spain | 1 | 1 |
| West Romania | 4 | 2 |

**Table S7:** Ploidy estimation results and FDSTools ploidy parameters. Available as Standalone.

The table is available as a comma separated value CSV file at https://doi.org/10.5281/zenodo.14686589. This data frame contains 1710 rows and 9 columns.
The columns are:

- **Sample_ID**: Name of the sample.
- **RandomForest_prob_2x**: Probability to belong to the diploid class according the Random Forest model trained in the study.
- **RandomForest_prob_4x**: Probability to belong to the tetraploid class (*i.e.* triploid or tetraploid) according the Random Forest model trained in the study.
- **RandomForest_prob_6x**: Probability to belong to the hexaploid class (*i.e.* pentaploid or hexaploid) according the Random Forest model trained in the study.
- **RandomForest_Prediction**: Predicted ploidy class.
- **Known_ploidy_class**: Known ploidy class of the reference samples.
- **Sample_class**: Indicates which samples are taken as reference to train the model and which samples were discarded at this step.
- **FDSTools_ploidy_parameter**: FDSTools Ploidy parameters used for the allele calling.
- **Reference_source**: Source of the ploidy for the reference samples.

**Table S8:** Quality and polymorphism of the 29 loci, assessed on the *Rosa gallica* dataset.

N_WAI: Number of alleles when considering the Whole Amplicon Information. N_length: Number of alleles when considering only allele length. N_rare: Number of alleles, detected by considering the Whole Amplicon information, with a frequency lower than 1%. PIC: Polymorphism Information Content.

| Loci | Calling strategy | Missing rate | N_WAI | N_length | N_rare | PIC |
|---|---|---|---|---|---|---|
| **Gal01_03** | RepeatFocused | 1.54 | 23 | 15 | 19 | 0.40 |
| **Gal02_11** | FullLength | 5.59 | 49 | 28 | 38 | 0.88 |
| **Gal02_14** | FullLength | 5.39 | 75 | 31 | 61 | 0.93 |
| **Gal03_19** | FullLength | 1.93 | 30 | 26 | 21 | 0.82 |
| **Gal03_25** | RepeatFocused | 0 | 14 | 9 | 9 | 0.52 |
| **Gal04_34** | RepeatFocused | 0.19 | 18 | 14 | 10 | 0.68 |
| **Gal05_38** | FullLength | 8.48 | 32 | 16 | 23 | 0.61 |
| **Gal06_42** | RepeatFocused | 12.52 | 38 | 25 | 27 | 0.86 |
| **Gal07_48** | RepeatFocused | 0.77 | 27 | 20 | 13 | 0.87 |
| **Gal07_51** | RepeatFocused | 0 | 11 | 8 | 7 | 0.61 |
| **Gal07_55** | FullLength | 2.12 | 51 | 21 | 44 | 0.71 |
| **Gal07_56** | FullLength | 5.01 | 36 | 16 | 22 | 0.80 |
| **Lab01-H9B07_07** | RepeatFocused | 0.58 | 10 | 7 | 3 | 0.75 |
| **Lab03-Rh58_24** | FullLength | 0.58 | 7 | 3 | 4 | 0.46 |
| **Lab07-Rw5G14_54** | RepeatFocused | 0 | 24 | 15 | 14 | 0.84 |
| **Trf01_04** | RepeatFocused | 0 | 13 | 8 | 6 | 0.75 |
| **Trf01_05** | RepeatFocused | 0 | 5 | 5 | 3 | 0.38 |
| **Trf02_08** | RepeatFocused | 0.58 | 32 | 22 | 18 | 0.86 |
| **Trf03_21** | RepeatFocused | 0 | 19 | 10 | 9 | 0.80 |
| **Trf03_23** | FullLength | 0 | 15 | 13 | 9 | 0.45 |
| **Trf04_27** | FullLength | 12.72 | 39 | 32 | 22 | 0.91 |
| **Trf04_30** | RepeatFocused | 5.59 | 27 | 14 | 18 | 0.83 |
| **Trf05_35** | FullLength | 7.51 | 32 | 20 | 19 | 0.80 |
| **Trf05_39** | RepeatFocused | 0 | 16 | 15 | 8 | 0.66 |
| **Trf06_44** | RepeatFocused | 10.4 | 59 | 23 | 38 | 0.93 |
| **Trf06_47** | FullLength | 0.39 | 40 | 15 | 26 | 0.79 |
| **Trf07_50** | RepeatFocused | 4.82 | 23 | 18 | 13 | 0.78 |
| **Trf07_52** | RepeatFocused | 4.82 | 76 | 34 | 58 | 0.93 |
| **Trf07_53** | RepeatFocused | 1.16 | 43 | 18 | 30 | 0.87 |
| **Average** | | 3.20 | 30.48 | 17.28 | 20.41 | 0.74 |

**Table S9:** DAPC cluster membership probabilities of the *R. gallica* dataset samples at $K = 12$ for the major mode. Available as Standalone.

The table is available as a comma separated value CSV file at https://doi.org/10.5281/zenodo.14686589. This data frame contains 867 rows and 15 columns.
The columns are:

- **Sample_ID**: Name of the sample.
- **Genetic_group_1** to **Genetic_group_12**: Membership probabilities for each given group.
- **Probability_be_Rosa_gallica**: Sum of the membership probabilities for the group 4, 5, 9 and 11 (See appendix D in main text).
- **Probability_not_Rosa_gallica**: Sum of the membership probabilities for the other groups (See appendix D in main text).

**Table S10:** Pairwise $\rho$ differentiation between geographical regions.

| Region | Slovenia/Croatia | Central Europe | Central France | Central East France | North East France | South East France | South West France | Germany | North Hungary | North Italy | West Poland | Poland/Ukraine | East Romania | North Romania | Ukraine Crimea |
|---|---|---|---|---|---|---|---|---|---|---|---|---|---|---|---|
| **Slovenia/Croatia** | 0 | 0.036 | 0.142 | 0.087 | 0.137 | 0.039 | 0.165 | 0.058 | 0.068 | 0.039 | 0.052 | 0.045 | 0.075 | 0.06 | 0.145 |
| **Central Europe** | 0.036 | 0 | 0.15 | 0.11 | 0.171 | 0.061 | 0.175 | 0.039 | 0.04 | 0.061 | 0.03 | 0.028 | 0.086 | 0.031 | 0.179 |
| **Central France** | 0.142 | 0.15 | 0 | 0.095 | 0.266 | 0.16 | 0.076 | 0.167 | 0.161 | 0.171 | 0.156 | 0.133 | 0.244 | 0.148 | 0.254 |
| **Central East France** | 0.087 | 0.11 | 0.095 | 0 | 0.189 | 0.093 | 0.113 | 0.118 | 0.112 | 0.074 | 0.121 | 0.09 | 0.155 | 0.105 | 0.188 |
| **North East France** | 0.137 | 0.171 | 0.266 | 0.189 | 0 | 0.153 | 0.308 | 0.146 | 0.178 | 0.182 | 0.153 | 0.158 | 0.231 | 0.2 | 0.29 |
| **South East France** | 0.039 | 0.061 | 0.16 | 0.093 | 0.153 | 0 | 0.182 | 0.053 | 0.105 | 0.033 | 0.089 | 0.054 | 0.127 | 0.064 | 0.19 |
| **South West France** | 0.165 | 0.175 | 0.076 | 0.113 | 0.308 | 0.182 | 0 | 0.2 | 0.195 | 0.195 | 0.197 | 0.158 | 0.241 | 0.174 | 0.264 |
| **Germany** | 0.058 | 0.039 | 0.167 | 0.118 | 0.146 | 0.053 | 0.2 | 0 | 0.065 | 0.076 | 0.064 | 0.034 | 0.115 | 0.034 | 0.198 |
| **North Hungary** | 0.068 | 0.04 | 0.161 | 0.112 | 0.178 | 0.105 | 0.195 | 0.065 | 0 | 0.107 | 0.036 | 0.048 | 0.122 | 0.061 | 0.178 |
| **North Italy** | 0.039 | 0.061 | 0.171 | 0.074 | 0.182 | 0.033 | 0.195 | 0.076 | 0.107 | 0 | 0.095 | 0.062 | 0.119 | 0.07 | 0.176 |
| **West Poland** | 0.052 | 0.03 | 0.156 | 0.121 | 0.153 | 0.089 | 0.197 | 0.064 | 0.036 | 0.095 | 0 | 0.04 | 0.134 | 0.059 | 0.186 |
| **Poland/Ukraine** | 0.045 | 0.028 | 0.133 | 0.09 | 0.158 | 0.054 | 0.158 | 0.034 | 0.048 | 0.062 | 0.04 | 0 | 0.096 | 0.012 | 0.181 |
| **East Romania** | 0.075 | 0.086 | 0.244 | 0.155 | 0.231 | 0.127 | 0.241 | 0.115 | 0.122 | 0.119 | 0.134 | 0.096 | 0 | 0.112 | 0.143 |
| **North Romania** | 0.06 | 0.031 | 0.148 | 0.105 | 0.2 | 0.064 | 0.174 | 0.034 | 0.061 | 0.07 | 0.059 | 0.012 | 0.112 | 0 | 0.198 |
| **Ukraine Crimea** | 0.145 | 0.179 | 0.254 | 0.188 | 0.29 | 0.19 | 0.264 | 0.198 | 0.178 | 0.176 | 0.186 | 0.181 | 0.143 | 0.198 | 0 |

**Table S11:** Private allelic richness and standard deviation per geographical region.

| Regional Group | Private Allelic Richness | StDev | Allelic Richness | StDev |
|---|---|---|---|---|
| Slovenia/Croatia | 0.39 | 0.10 | 5.82 | 0.45 |
| Central Europe | 0.24 | 0.07 | 5.33 | 0.47 |
| Central France | 0.22 | 0.06 | 4.68 | 0.38 |
| Central East France | 0.48 | 0.18 | 5.48 | 0.39 |
| North East France | 0.24 | 0.08 | 4.69 | 0.44 |
| South East France | 0.23 | 0.10 | 5.34 | 0.43 |
| South West France | 0.24 | 0.07 | 4.66 | 0.32 |
| Germany | 0.25 | 0.07 | 5.36 | 0.47 |
| North Hungary | 0.27 | 0.08 | 5.45 | 0.49 |
| North Italy | 0.56 | 0.12 | 6.06 | 0.50 |
| West Poland | 0.15 | 0.05 | 4.89 | 0.42 |
| Poland/Ukraine | 0.25 | 0.05 | 5.38 | 0.46 |
| East Romania | 0.36 | 0.15 | 4.60 | 0.39 |
| North Romania | 0.30 | 0.09 | 5.17 | 0.45 |
| Ukraine Crimea | 0.94 | 0.21 | 5.21 | 0.42 |

### 4 Codes and Scripts description

Following lines describe content of the **rosa_gallica_genpop** folder.

#### Folder structure

##### Root file

- **rosa_gallica_genpop.Rproj**
  RStudio project file.

##### programs/

Analysis scripts written in Quarto (`.qmd`) and R.

- **1_Analysis_test_param_FDSTools.qmd**
  Testing and comparison of FDSTools parameter sets for each locus.

- **2_Paralogue_detection.qmd**
  Identification of potential paralogous loci.

- **3_Ploidy_estimation.qmd**
  Estimation of individual ploidy levels using random forest.

- **4_FDSTools_final_calling.qmd**
  Final allele calling using selected FDSTools parameters.

- **5_Allele_dosage_part1_allele_competition_investigation.qmd**
  Investigation of allele competition patterns.

- **5_Allele_dosage_part2_Tetraploids.qmd**
  Allele dosage inference for tetraploid individuals.

- **5_Allele_dosage_part3_Hexaploids.qmd**
  Allele dosage inference for hexaploid individuals.

- **5_Allele_dosage_part4_file_merging.qmd**
  Merging allele dosage outputs across ploidy levels.

- **6_Missing_data_optimization.qmd**
  Optimization and filtering based on missing data.

- **7_Clone_detection_refined.qmd**
  Detection and filtering of clonal individuals.

- **8_Amplicon_demultiplexing.qmd**
  Demultiplexing and preprocessing of amplicon sequencing data.

- **9_kmer_based_dist_computation.qmd**
  Computation of k-mer–based genetic distances.

- **10_Mis_ident_detection_refined.qmd**
  Detection of potential sample misidentifications.

- **11_Marker_description_Gallica.qmd**
  Description and summary of retained markers.

- **12_Gallica_structure_refined.qmd**
  Population genetic analyses.

- **Custom_functions.R**
  Custom R functions used throughout the analyses.

**data/**

Raw data, intermediate files, and formatted outputs.

- **Amplicon_sorting/**
  FDSTools input files and locus parameter definitions used just before amplicon demultiplexing.

- **Calling_diplo/**, **Calling_tetra/**, **Calling_hexa/**
  Allele calling results for diploid, tetraploid, and hexaploid individuals.

- **calling_all/**
  Combined allele calling outputs across all ploidy levels, including GenAlEx-formatted files, final genotypic tables, and allele information files.

- **calling_all/allele_loci_fasta/**
  FASTA files of allele sequences per locus and corresponding distance matrices. This folder will be filled while running the pipeline.

- **Genotypic_table_all_ploidy_detection/**
  Files used for ploidy detection and validation, including diagnostic loci, metadata, and final ploidy assignments.
- **Genotypic_table_pilot_run_FDSTools_param/**
  Pilot genotypic tables generated under multiple FDSTools parameter combinations.
- **Genotypic_table_pilot_run_paralog_detection/**
  Pilot datasets used for paralogue detection.
- **Rgall_structure/**
  Input and output files for population structure and diversity analyses (e.g. ADZE, Genodive, STRUCTURE/KFinder).
- **Metadata_genotyped.txt**
  Metadata associated with genotyped individuals.
- **fixed_ind_code.txt**
  Table of correspondances between individual identifiers.

**results/**

Processed results and summary outputs used for downstream analyses and figures.

- **AMOVA_region/**
  AMOVA analyses by region, including Genodive inputs and outputs.
- **Genotypic_table_all_ploidy_detection/**
  Final ploidy assignments used in downstream analyses.
- **Rgall_structure/**
  Outputs related to population structure and null allele analyses.
- **SSRseq_pipeline_param_choice/**
  Summary of the selected SSRseq and FDSTools parameters.

**Notes**

- File names and directory structure reflect the analytical workflow.
- Scripts are numbered to indicate their logical execution order.
- Some intermediate files are not available here but can be easily regenerated using the qmd and R files located in the **programs** folders.
